# A bivalent VP2-based vaccine approach reveals co-expression-driven interference between VP2 antigens impairing BTV-neutralizing antibody induction

**DOI:** 10.64898/2026.11.11.764333

**Authors:** Luis Jiménez-Cabello, Juana Sánchez-Puig, Iván Mazuecos-Aragonés, Dunia Asensio-Cob, Julio Benavides, Sergio Utrilla-Trigo, María M. Lorenzo, Aitor Nogales, Eva Calvo-Pinilla, Rafael Blasco, Javier Ortego

## Abstract

Bluetongue virus (BTV) is an important arboviral pathogen of ruminants that causes a significant impact on the primary sector. More than 29 different BTV serotypes exist, hindering control and prevention of Bluetongue (BT) disease. Conventional vaccines possess safety and efficacy limitation and do not permit to confer multiserotype protection and differentiate between vaccinated and infected animals (DIVA strategy). Here, we developed recombinant Modified Vaccinia Ankara (rMVA) viral vectors individually expressing or co-expressing the neutralizing antibody (nAb) target proteins VP2 of BTV serotypes 4 and 8. We evaluated their immunogenicity and protective efficacy in IFNAR(-/-) mice and sheep. Immunization of IFNAR(-/-) mice with the rMVA co-expressing of BTV-4 VP2 and BTV-8 VP2 led to abolished nAb induction while eliciting non-neutralizing antibodies against both VP2 proteins, suggesting a previously unrecognized interference between VP2 antigens. Combined immunization of IFNAR(-/-) mice with rMVAs expressing either BTV-4 VP2 or BTV-8 VP2 elicited a strong neutralizing immune response and complete protection against lethal challenge with BTV-4 or BTV-8. In sheep, combinatorial prime-boost immunization with these monovalent rMVAs induced nAbs against both homologous BTV serotypes. Immunized sheep were fully protected against BTV-4, as no fever or viral replication were detected after challenge. Similarly, immunized sheep were also protected against BTV-8 although detectable but mild RNAemia and slight increase in rectal temperatures were observed after challenge. Overall, we identified a relevant constraint for multivalent VP2-based vaccine design and present a DIVA vaccination strategy that offers bivalent protection against BTV in a natural host of the disease.

**IMPORTANCE:** Emergence and co-circulation of virulent strains of BTV pose one of the most significant challenges for animal health and the primary sector. Available vaccines against BTV have been effective tools to counteract BT but they confer serotype-specific protection and do not permit to establish a DIVA strategy. Besides, recently commercialized inactivated vaccines against the highly virulent BTV-3 circulating in Europe do not offer adequate protection in livestock. Therefore, advancing next-generation vaccine strategies has become a critical priority. In this work, we present a novel DIVA vaccination strategy based on the recombinant MVA vaccine vector. We engineered recombinant MVAs expressing the structural proteins VP2 of serotype 4 or 8 and confirmed their potential to confer robust bivalent protection in a natural BTV host species. Although this new vaccination regime is not expected to provide broad-spectrum protection, it solves safety concerns of classic vaccine approaches, represents a DIVA-compatible vaccination strategy and can be readily tailored to confront multiserotype BTV epizootics. In addition, our work reveals a critical constraint for next-generation vaccine design against BTV and related orbiviruses, providing important insights for the future guidance to design novel and effective vaccines against orbiviruses regarding the formulation of the VP2 antigen.

## INTRODUCTION

Bluetongue (BT), caused by Bluetongue virus (BTV), is a Culicoides-transmitted disease with a significant impact on animal health, causing severe economic losses due to its worldwide distribution and the wide range of affected hosts ^1–3^. Sheep and some wild ruminants like white-tailed deer (*Odocoileus virginianus*) are the most affected hosts, showing considerable mortality rates, whereas cattle, goats and most wild ruminants are usually asymptomatic. Nevertheless, cattle may show signs of BT disease _4–8_. BTV circulates in tropical and subtropical regions between 50° N and 35° S. Nonetheless, new incursions have been detected in Europe every year since 1998 ^9,10^. Indeed, recent and important outbreaks of BTV-3 and BTV-12 have occurred in Europe ^11–13^.

BTV (genus *Orbivirus*, family *Sedoreoviridae*) is characterized by an icosahedral capsid (∼90 nm in diameter) divided into three concentric protein layers that harbor a segmented dsRNA viral genome _14–16_. The outer capsid of the virus consists of 180 copies (60 trimers) of protein VP2 together with protein VP5, which is also involved in virus entry ^17,18^. The inner capsid is composed of the subcore, formed by protein VP3, and the intermediate layer, constituted by protein VP7, along with three structural proteins with enzymatic activities: VP1, VP4 and VP6 ^19–21^. Additionally, the viral genome encodes five nonstructural proteins (NS1, NS2, NS3/NS3A, NS4 and NS5) ^22,23^.

More than 29 serotypes of BTV have been described through phylogenetic studies, sequencing data and cross-neutralization assays ^24^. The main determinant of serotype specificity is the outer capsid protein VP2, the most exposed virion protein ^24^. This protein plays a pivotal role during the initial phases of viral infection ^25,26^. The trimers of protein VP2, which contains sialic acid-binding domains ^27,28^, bind to the cellular receptor enabling viral adhesion and entry into the host cell via clathrin-dependent endocytosis ^29,30^. Among the different structural BTV proteins, protein VP2 is the main target of neutralizing antibodies (nAbs) ^31,32^. Although cross-reactivity between proteins VP2 of different serotypes exists, nAbs induced by this protein are mainly serotype-specific and confer protection against homologous BTV serotypes ^33,34^. In consequence, the absence of cross-neutralization between different BTV serotypes prevails, varying among ruminant species and occurring mostly in laboratory animal models ^35^.

Due to the lack of efficacious therapeutic treatments ^36–39^, vaccination is the most effective countermeasure for the successful control of BTV ^40^. Classic vaccine approaches have been implemented for BT control. BTV live attenuated vaccines (LAVs) are highly immunogenic and inexpensive, but are related with animal welfare and productivity drawbacks, potential transmission to the insect vector, reversion to virulence or genomic reassortment with field strains ^41–46^. Despite inactivated vaccines are safer and permit to control BTV, they are less immunogenic and more expensive than LAVs ^47–50^. Importantly, both conventional vaccination approaches against BTV only induce serotype-specific protection and do not allow differentiation between naturally infected and vaccinated animals (DIVA character) ^50–52^, which motivates the development of next-generation vaccines against BTV.

An ideal BTV vaccine should be affordable, completely safe, DIVA and capable of inducing multiserotype and long-lasting protection in all ruminant species ^50^. To date, few experimental vaccine candidates possess multiserotype protective potential ^50–52^. Different BTV antigens have been evaluated as targets for multiserotype vaccine development such as the highly conserved non-structural proteins NS1 and NS2 that harbor T CD8+ epitopes within their sequence ^53–57^ and are related with faster viral clearance and milder pathology induced by BTV infection ^54,55,58^. Lately, we observed that the combined expression of NS1 and NS2-Nt with protein VP2 of BTV-4 confers full protection against the homologous BTV-4 but detectable virus replication is still observed after heterologous challenge with BTV-8 in natural hosts ^59,60^. Different studies have also pointed out the highly conserved BTV protein VP7 as a inducer of multiserotype protective cell-mediated immune responses against BTV ^61–63^. However, the role of orbiviral protein VP7 as an appropriate antigen for vaccine development must be clarified ^59,64^. Overall, experimental vaccine candidates based on proteins VP7, NS1 and/or NS2 are usually related with significant levels of viral replication and, in some cases, with BTV-associated pathology. Therefore, the inclusion of VP2 in the vaccine composition remains the most promising and rational approach for achieving safe, DIVA-compatible and complete multiserotype protection against the wide serotype diversity of BTV.

As protein VP2 is the target of nAbs and induction of a neutralizing response against BTV correlates with protection, a wider range of recombinant vaccine candidates based on protein VP2 have been developed ^40,50–52^. However, protective nAbs responses induced by expression or delivery of protein VP2 are serotype-specific ^51,65^ like those induced by classic approaches ^32^. Therefore, experimental vaccine approaches based on neutralizing responses against BTV that aims to confer broad-spectrum protection require the formulation of VP2 proteins of different serotypes. This is the case of Virus-like particles (VLPs) (composed of proteins VP2, VP5, VP7 and VP3), which are immunogenic and protective in natural hosts when adjuvanted ^33,66^, but multiserotype protection depends on immunization with cocktails of VLPs from different serotypes ^67–69^. Similarly, LAVs artificially engineered by suppressed expression of proteins VP6/NS4 or NS3/NS3A, and named DISC ^70^ or DISA ^71^ vaccines, respectively, are safe and effective in natural hosts against homologous BTV serotypes but multiserotype protective responses are conditioned by formulation of a plethora of viruses of different serotypes ^72–74^.

In this work, we evaluate the protection against BTV serotypes 4 and 8 conferred by different vaccination strategies based on the MVA viral vaccine vector, which expresses the VP2 proteins of these serotypes either individually or in combination. We engineered rMVA viral vectors individually expressing or co-expressing serotype 4 VP2 and serotype 8 VP2 of BTV. After characterizing the generated rMVA viral vectors expressing these BTV antigens, we evaluated their immunogenicity, and the homologous protection induced against BTV in BTV-susceptible IFNAR(-/-) mice. Thereafter, the protective efficacy against BTV of a homologous prime-boost immunization regime combining rMVAs individually expressing serotype 4 VP2 and serotype 8 VP2 of BTV in the same inoculum was evaluated in sheep, one of the most affected natural hosts of BTV.

## MATERIALS AND METHODS

### Cells lines and viruses

Chicken embryo fibroblasts (DF-1) (ATCC, Cat. No. CRL-12203) were grown in Dulbecco’s Modified Eagle’s medium (DMEM) (Biowest, Nuaillé, France) supplemented with 2mM glutamine (Gibco, Waltham, MA, USA) and 10% fetal calf serum (FCS) (Gibco, Waltham, MA, USA). Green monkey kidney cells (Vero) (ATCC, Cat. No. CCL-81) were grown in DMEM supplemented with 2mM glutamine and 5% FCS.

BTV serotype 1 (ALG2006/01) (BTV-1), BTV serotype 4 Morocco strain (MOR2009/09) (BTV-4M), BTV serotype 4 (SPA2004/02) and BTV serotype 8 (BEL/2006) (BTV-8) were used in the experiments. BTV-4M strain is a reassortant strain between BTV-1 (segments 1, 4, 5, 7, 9, 10) and BTV-4 (segments 2, 3, 6, 8) isolated from sheep blood in KC insect cells _75,76_. Virus stocks and titrations were performed by standard methods previously described^77^.

### Generation of rMVA vaccine vectors

Recombinant modified vaccinia Ankara (rMVA) viral vectors were engineered to express BTV VP2 proteins from serotypes 1 (SPA2008/01; GenBank Accession Number: KP821020.1), 4 (SPA2004/02; GenBank Accession Number: KP821068.1) and 8 (NET2006/04; GenBank Accession Number: JX680448.1) either individually or combined. The VP2 genes from the different BTV serotypes were synthesized by GenScript and cloned in an MVA transfer plasmid containing the vaccinia virus A27L or F13L genes and the corresponding flanking sequences for homologous recombination.

Recombinant MVA viruses MVA-VP2_A4_ and MVA-VP2_A8_ were obtained by homologous recombination of the transfer plasmids with a mutant MVA virus lacking gene A27L (MVA-ΔA27L) following protocols already described _78_. For this, BHK-21 cells were infected with MVA-ΔA27L at a MOI of 0.05 PFU/cell. Infected cells were subsequently transfected with 2 µg of the corresponding plasmids (pMVA-A-BTV-VP2-4 y pMVA-A-BTV-VP2-8) using 6 µl of Fugene HD and incubated for 3 days. Recombinant viruses were clonally selected from the progeny by three successive rounds of plaque purification in agarose, selecting plaques that had recovered normal size and showed loss of fluorescence.

Double recombinant viruses MVA-VP2_A4_-VP2_F8_ and MVA-VP2_A4_-VP2_F1_, were obtained by inserting the corresponding VP2 gene downstream of the F13L gene into the previously obtained MVA-VP2_A4_ and MVA-VP2_A8_ viruses. With this aim, the F13L gene was first deleted in those viruses by a modification of the Cre-LoxP strategy used previously for A27L deletion ^78^. First, an intermediate MVA-A-BTV-VP2_A4_ virus containing a floxed F13L gene was isolated by recombination with plasmid pMVA-LfoptLRed, containing a floxed, codon modified F13 gene as well as gene for fluorescent protein dsRed, between the F13L recombination flanks. Virus MVA-VP2_A4_-ΔF13L was isolated by sequential passaging of the virus in BHK-21 expressing Cre recombinase until no viruses producing large plaques were detectable. Finally, virus MVA-VP2_A4_-ΔF13L was used to insert a second VP2 gene by recombination with plasmids pMVA-F-BTV-VP2-1 or pMVA-F-BTV-VP2-8 containing the F13L gene and VP2 from BTV serotype 1 or serotype 8 placed between F13L recombination flanks. Recombinant viruses were identified and isolated based on the reconstitution of normal plaque size ^79^. This strategy resulted in the generation of dual-expression recombinant viruses MVA-VP2_A4_-VP2_F8_ and MVA-VP2_A4_-VP2_F1_.

### Indirect immunofluorescence microscopy

DF-1 cells were grown in glass coverslips and infected with parental MVA, MVA-VP2_A4_, MVA-VP2_A8_ or MVA-VP2_A4_-VP2_F8_ at a MOI of 0.1, or non-infected. Twenty-four hours after infection, cell monolayers were fixed for 15 minutes with 4% paraformaldehyde. Fixed cells were blocked with 20% FBS-PBS-Saponine 0.2% (20% blocking solution) for 60 min at room temperature (RT). DF-1 cells were incubated overnight at 4°C with a mouse serum against recombinant protein VP2 of BTV-4 or BTV-8 (1:500) diluted in PBS-FBS. After three serial washing steps with PBS, DF-1 cells were incubated for 30 min at room temperature (RT) with Alexa Fluor 488 goat conjugated anti-mouse IgG (Invitrogen™, German Town, MD, USA) (1:1000). Coverslips with infected or non-infected DF-1 cells were washed three times with PBS and once with PBS-DAPI (1:10,000), and visualized in a Zeiss Axio fluorescence microscope (Zeiss, Oberkochen, Germany). Adobe Photoshop CS5 Extended (Adobe Systems, CA, USA) was used afterwards for image editing.

Regarding Supplementary data, DF-1 cells were grown in glass coverslips and infected with parental MVA or MVA-VP2_A4_-VP2_F1_ at a MOI of 1, or non-infected. Twenty-four hours after infection, cell monolayers were fixed for 15 minutes with 4% paraformaldehyde. Fixed cells were blocked with 20% FBS-PBS-Saponine 0.2% (20% blocking solution) for 60 min at room temperature (RT). DF-1 cells were incubated overnight at 4°C with a mouse serum against recombinant protein VP2 of BTV-4 or BTV-1 (1:500) diluted in PBS-FBS. After three serial washing steps with PBS, DF-1 cells were incubated for 30 min at room temperature (RT) with Alexa Fluor 488 goat conjugated anti-mouse IgG (Invitrogen™, German Town, MD, USA) (1:1000). Coverslips with infected or non-infected DF-1 cells were washed three times with PBS and once with PBS-DAPI (1:10,000), and visualized in a Zeiss Axio fluorescence microscope (Zeiss, Oberkochen, Germany). Adobe Photoshop CS5 Extended (Adobe Systems, CA, USA) was used afterwards for image editing.

### Analysis of rMVA virus growth

Monolayers of permissive DF-1 cells were infected with parental MVA, MVA-VP2_A4_, MVA-VP2_A8_ or MVA-VP2_A4_-VP2_F8_ at a MOI of 0.1. At 24, 48, and 72 h.p.i., cells were harvested, and virus titers in cell lysates were determined by a plaque assay in DF-1 cells, as previously described^80^.

### Mice and sheep

Type I interferon receptor defective mice (IFNAR (-/-)) on a 129 Sv/Ev background and sheep (Spanish “Churra” sheep breed) were used for the studies. All mice and sheep used were matched for age (8 weeks and 6 months, respectively). Mice and sheep were housed under pathogen-free conditions and allowed to acclimatize to the biosafety level 3 (BSL3) animal facilities at the Animal Health Research Center (INIA-CISA), Madrid, before use.

### Mice immunization and challenge

To assess the immunogenicity of the rMVAs, two groups of IFNAR(-/-) mice (n=4) were intramuscularly immunized following a homologous prime-boost regime consisting of two doses of 1x10^7^ PFU per mouse of MVA-VP2_A4_, MVA-VP2_A8_ or MVA-VP2_A4_-VP2_F8_, administered four weeks apart. Another group of IFNAR(-/-) mice (n=4) was intramuscularly immunized following a homologous prime-boost regime consisting of two doses (administered in a four-weeks interval) of 1x10^7^ PFU per mouse of MVA-VP2_A4_ and MVA-VP2_A8_ conducted separately in different sites of inoculation (different legs). An additional group of IFNAR(-/-) mice (n=4) was intramuscularly immunized following a homologous prime-boost regime consisting of two doses (administered in a four-weeks interval) of 1x10^7^ PFU per mouse of each MVA-VP2_A4_ and MVA-VP2_A8_ mixed in the same inoculum. A group of IFNAR(-/-) mice (n=4) was left untreated (control). Submandibular blood collection was carried out at 4 w.p.p. and 3 w.p.b. to analyze the humoral immune response.

To evaluate the protective potential of the rMVAs, two groups of IFNAR(-/-) mice (n=5) were intramuscularly immunized following a homologous prime-boost regime consisting of two doses of 1x10_7_ PFU per mouse of each MVA-VP2_A4_ and MVA-VP2_A8_ (inoculated separately in different legs), administered four weeks apart. A group of IFNAR(-/-) mice (n=5) was left untreated (control) for each challenge. Animals were subcutaneously challenged with a lethal dose of BTV-4M (10 PFU) or BTV-8 (100 PFU) at 3 w.p.b. In all cases, submandibular blood collection was carried out in mice after virus challenge at 3, 5, 7, 10 and 14 d.p.i. for the analysis of RNAemia.

### Sheep immunization and challenge

16 naive healthy sheep (Spanish “Churra” sheep breed), aged 6 months, were acclimated for seven days at the BSL3 animal facility of the Animal Health Research Center (INIA-CISA) before starting the experiment. All sheep involved in the experiment were negative to BTV by ELISA. Briefly, two groups of sheep (n=4) were intramuscularly immunized following a homologous primer-boost strategy consisting of two doses of 10^8^ PFU of each MVA-VP2_A4_ and MVA-VP2_A8_ (inoculated separately in different legs), administered four weeks apart. Two groups of sheep were untreated (control). Pre-challenge blood samples were collected from all animals at 3 w.p.b. Non-immunized and immunized sheep were subcutaneously challenged with a dose of 10_5_ PFU of BTV-4M or BTV-8 at 3 w.p.b. After virus challenge, blood collection for virological analyses was conducted by specialized veterinary personnel at 3, 5, 7, 10, 12, 14, and 18 d.p.i. Rectal temperature measurements were conducted every day from four days prior to challenge until 18 d.p.i. Fever threshold was determined for each group as the mean rectal temperature plus three times the standard deviation of the temperatures recorded for four days before challenge. On day 18 post-infection all sheep were euthanized.

Regarding Supplementary data, 8 naive healthy sheep (Spanish “Churra” sheep breed), aged 6 months, were acclimated for seven days at the BSL3 animal facility of the Animal Health Research Center (INIA-CISA) before starting the experiment. All sheep involved in the experiment were negative to BTV by ELISA. A group of sheep (n=4) was intramuscularly immunized following a homologous primer-boost strategy consisting of two doses of 10_8_ PFU of MVA-VP2_A4_-VP2_F1_, administered four weeks apart. A group of sheep was untreated (control). Pre-challenge blood samples were collected from all animals at 3 w.p.b. Non-immunized and immunized sheep were subcutaneously challenged with a dose of 10_5_ PFU of BTV-4M at 3 w.p.b. After virus challenge, blood collection for virological analyses was conducted by specialized veterinary personnel at 3, 5, 7, 10, 12, 14, and 18 d.p.i. Rectal temperature measurements were conducted every day from four days prior to challenge until 18 d.p.i. Fever threshold was determined for each group as the mean rectal temperature plus three times the standard deviation of the temperatures recorded for four days before challenge. On day 18 post-infection all sheep were euthanized.

### Viraemia and RNAemia analysis by plaque assay and RT-qPCR

Blood samples were collected at 3, 5, 7, 10 and 14 d.p.i from the submandibular plexus of mice and at 3, 5, 7, 10, 12, 14, and 18 d.p.i. from sheep with EDTA as anti-coagulant.

For the analysis of RNAemia by RT-qPCR, RNA was extracted from 50 µL of blood using TRIzol Reagent (Sigma Aldrich, St. Louis, MO, USA) following the protocol established by the manufacturer. Viremia was analyzed in duplicate by real-time RT-qPCR specific for BTV segment 5 (encoding for NS1). The real-time RT-qPCR specific for BTV segment 5 was performed using primers and probe described by Toussaint et al. ^81^. Only Ct values lower than 38 were considered indicative of RNAemia (positive), according to the cut-off established by Toussaint et al. ^81^.

For the analysis of viraemia by plaque assay, 50 µL of sheep blood were diluted in PBS1X and centrifuged at 3000 rpm for 10 minutes. Thereafter, the supernatant was removed, and the pellet was lysed in 450 µL of sterile water for 2 minutes. Cell lysis was stopped by adding 50 µL of PBS10X. Then, samples were inoculated into 12-well plates containing semi-confluent monolayers of Vero cells. Following incubation for 1 h, an agar overlay (DMEM-10%-FBS-0.4%-Noble Agar, Becton Dickinson, MD, USA) was added and plates were incubated for 5 days at 37°C in 5% CO_2_. Plaques were fixed with 10% formaldehyde and visualized with 2% crystal violet-PBS.

### Blood measurements

A multiparameter autohematology analyzer (BC-5300 Vet; Mindray, China) was used to determine the total and differential cell counts in sheep blood for each group and collected into EDTA tubes.

### Determination of circulating levels of cytokines

Sera from immunized and non-immunized sheep were collected the day before the challenge and at 3 and 5 d.p.i. Circulating cytokine levels were analyzed using a multiplex fluorescent bead immunoassay for quantitative detection of sheep cytokines (MILLIPLEX® Ovine Cytokine/Chemokine Panel 1, Burlington, MA, USA). Samples were analyzed for IL1-α, IL-1β, IL-6, IL-10, and TNF with a MAGPIX system (Luminex Corporation, Austin, TX, USA). Values of blank samples were subtracted from values of samples.

### Specific Indirect ELISA for Proteins VP2 of BTV-4 and BTV-8

MaxiSorp plates (Nunc) (Thermo Fisher Scientific, New York, NY, USA) were coated with either BTV-4 VP2 (200 ng per well) or BTV-8 VP2 (200 ng per well) purified baculovirus-expressed proteins in PBS and incubated overnight at 4°C. Plates were saturated with blocking buffer (PBS-0.05%-Tween 20-5% skim milk). Individual mice sera diluted in blocking buffer (1:200) were added and incubated for 2h at 37°C. After three washes with PBS-0.05% Tween 20, plates were incubated for 1 h at 37°C with an anti-mouse-IgG-HRP secondary antibody (BioRad, Hercules, CA, USA) (1:2000) in diluted blocking buffer (PBS-0.05%-Tween 20-1% skim milk). Finally, after three washes in PBS-0.05% Tween 20, the reaction was developed with 50µL of TMB (Thermo Fisher Scientific, Rockville, MD, USA) and stopped by adding 50µL of 4 N H_2_SO_4_ (Merck, Darmstadt, Germany). Results were expressed as optical densities (ODs) measured at 450 nm.

### Plaque reduction neutralization test (PRNT)

Two-fold dilutions (from 1:5) of heat inactivated mice or sheep sera (56°C for 30 minutes) were incubated with 100 PFU of BTV-4, BTV-8 or BTV-1, for 1h at 37°C. Then, samples were inoculated into 12-well plates containing semi-confluent monolayers of Vero cells. Following incubation for 1h, an agar overlay (DMEM-10%-FBS-0.4%-Noble Agar (Becton Dickinson, MD, USA)) was added and plates were incubated for 5 days at 37°C in 5% CO_2_. Plaques were fixed with 10% formaldehyde and visualized with 2% crystal violet-PBS. PRNT_50_ titer was calculated as the highest dilution of serum that neutralized 50% of the control virus input.

### Protein structure prediction and VP2-VP2 interface analysis

Three-dimensional (3D) structural models of VP2 from Bluetongue virus serotype 4 (BTV-4) and serotype 8 (BTV-8) were generated using AlphaFold 3 (AF3) through the AlphaFold Server web platform (https://alphafoldserver.com) using default settings. Predictions were performed for VP2 monomers as well as homologous and heterologous trimeric assemblies. For each representative, the highest-ranking structural model, determined by the predicted local distance difference test (pLDDT) score, was retained. Predicted structures were visualized and inspected using UCSF ChimeraX (version 1.12) (114). Trimeric assemblies were subsequently analyzed to compare interaction patterns between VP2 protomers in homologous and heterologous (mixed BTV-4/BTV-8) complexes. To characterize intermolecular contacts and interface properties, all trimeric models were submitted to the Protein Interfaces, Surfaces and Assemblies (PISA) server available through the Protein Data Bank in Europe (PDBePISA). PISA was used to perform a detailed analysis of VP2–VP2 interfaces, including the calculation of the number of interface residues, buried surface area (BSA), hydrogen bonds, salt bridges, solvation free-energy gain upon interface formation (ΔG), and the Complex Formation Significance Score (CSS).

### Statistical Analysis

Data were analyzed using GraphPad Prism version 8.0.1 (GraphPad Software, San Diego, CA, USA). Survival curves for each immunized mice group were compared to those of non-immunized mice in search of statistical differences using Log-rank test. Comparisons of mean responses between groups in the ELISA assays for mice groups were performed using one-way ANOVA (Kruskal-Wallis test). Comparisons of mean responses between groups in the viremia and RNAemia analysis for mice groups were performed using Mann-Whitney non-parametric test. Comparisons of mean responses between groups for the viral growth kinetics analysis, PRNT_50_ as well as data on sheep rectal temperature, viraemia, RNAemia, hematologic values and cytokine levels were conducted by two-way ANOVA with a post hoc Tukey test for multiple comparisons. A p-value lower than 0.05 was considered significant in all cases.

### Ethics statement

Animal experimental protocols were approved by the Ethical Review Committee at the INIA-CISA and Comunidad de Madrid (Permit number: PROEX 060.7/21) in strict accordance with EU guidelines 2010/63/UE about protection of animals used for experimentation, and other scientific purposes and Spanish Animal Welfare Act 32/2007.

## RESULTS

### Evaluation of BTV-4 VP2 and BTV-8 VP2 expression from rMVAs

To develop a bivalent MVA-based vaccine capable of simultaneously express nAbs-target VP2 antigens from two epidemiologically relevant BTV serotypes, we generated a rMVA viral vector expressing protein VP2 of BTV-4 along with protein VP2 of BTV-8 (MVA-VP2_A4_-VP2_F8_). Additionally, we designed rMVAs individually expressing these same proteins, VP2 of BTV-4 (MVA-VP2_A4_) and BTV-8 (MVA-VP2_A8_). VP2 genes were placed under the control of a synthetic Early/Late Poxviral promoter and inserted downstream of the A27L or F13L genes (Fig. 1A). Viruses were named according to the insertion place (A or F) and the number of the VP2 serotype.

**Figure 1.**
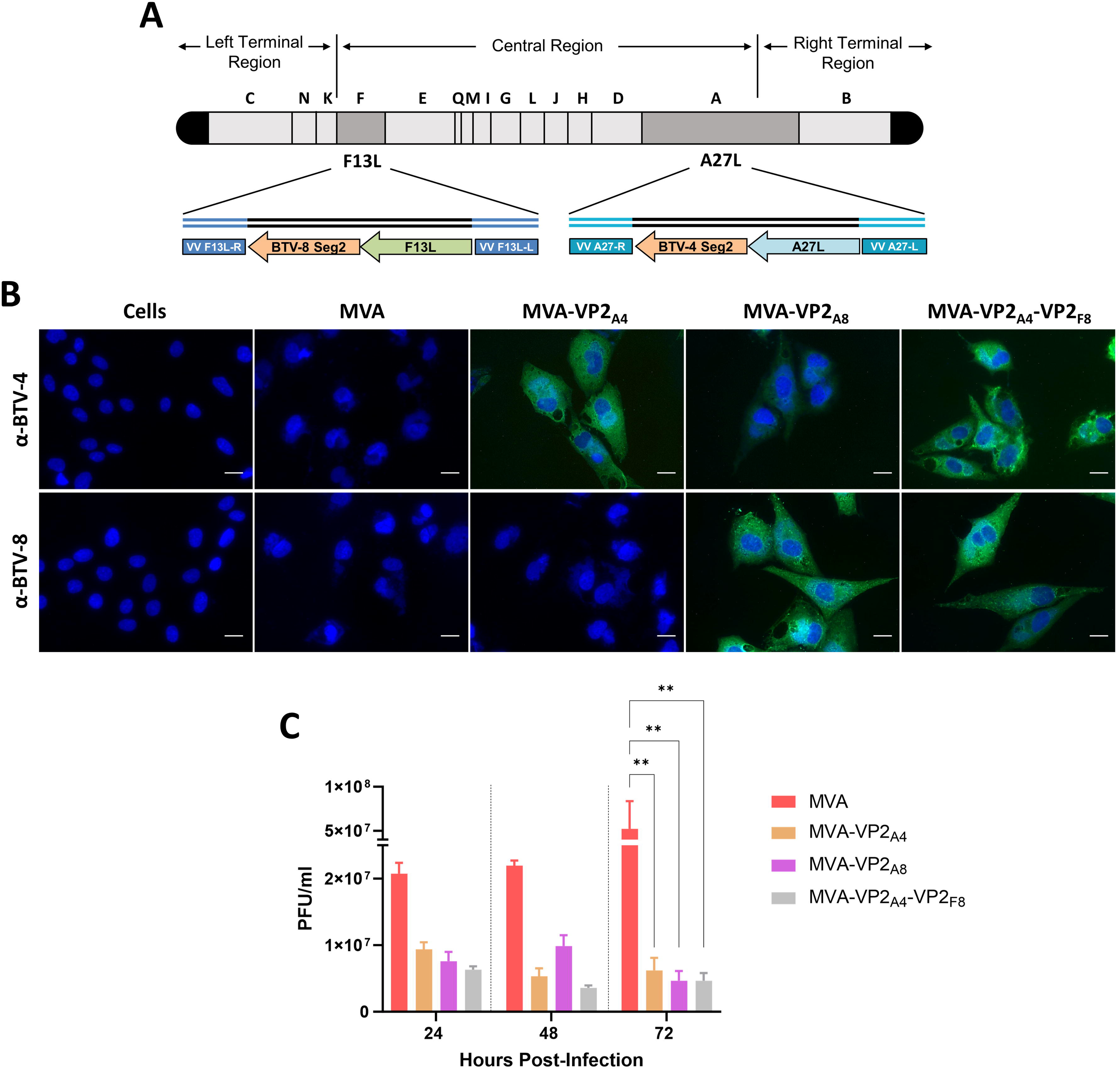
Expression analysis of heterologous BTV proteins by rMVA. (A) Schematic representation of the cloning strategy within the MVA genome. Segment 2 of BTV-8 and BTV-4 were cloned in the F13L and A27L loci, respectively. (B) Indirect immunofluorescence of DF-1 cells infected (MOI = 0.1) with parental MVA, MVA-VP2_A4_, MVA-VP2_A8_, MVA-VP2_A4_-VP2_F8_ or non-infected (control). Proteins VP2 of BTV-4 and VP2 of BTV-8 (green) were detected using mouse polyclonal sera against BTV-4 or BTV-8. Nuclei were stained with DAPI. Scale bars 20 µm. (C) Viral growth kinetics of parental MVA, MVA-VP2_A4_, MVA-VP2_A8_ and MVA-VP2_A4_-VP2_F8_. Monolayers of permissive DF-1 cells were infected with parental MVA, MVA-VP2_A4_, MVA-VP2_A8_ or MVA-VP2_A4_-VP2_F8_ (MOI = 0.1). At 24, 48 and 72 h.p.i., virus titers in cell lysates were quantified by plaque assay. The assay was performed in triplicate. Bars represent the mean PFU/mL value of each group and error bars represent SD. Statistical differences between replication curves of parental MVA, MVA-VP2_A4_, MVA-VP2_A8_ and MVA-VP2_A4_-VP2_F8_ were calculated by two-way ANOVA (post hoc Tukey test for multiple comparisons). ** P value < 0.002.

We first confirmed the proper expression of the heterologous BTV antigens cloned in the rMVAs by indirect immunofluorescence assay (IFA). We noticed the expression of proteins VP2 of BTV-4 and BTV-8 throughout the cytoplasm of DF-1 cells infected with MVA-VP2_A4_ and MVA-VP2_A8_, respectively (Fig. 1B).

Similarly, DF-1 cells infected with MVA-VP2_A4_-VP2_F8_ showed cytoplasmic fluorescent signals corresponding to the expression of both BTV genes (Fig. 1A). We did not observe evidence of a specific signal of BTV-4 VP2 or BTV-8 VP2 in non-infected or parental MVA (MVA-wt) infected cells (Fig. 1B). These results confirm the expression of both VP2 antigens by the rMVAs and suggest that their co-expression by a single rMVA does not noticeably disturb their expression.

In previous studies, we observed that cloning of protein VP2 of a highly related orbivirus (EHDV-8) mildly weakens MVA vector replication in DF-1 cells ^64^. Therefore, we analyzed the effect of cloning proteins VP2 of serotype 4 and 8 of BTV on MVA viral replication *in vitro*. To do so, we studied the growth kinetics of the generated rMVAs in DF-1 cells (multiplicity of infection (MOI) =0.1). Viral growth of MVA-VP2_A4_ and MVA-VP2_A8_ was very similar to that of parental MVA at 24 and 48 h.p.i. (Fig. 1C). Nonetheless, these rMVAs showed significantly lower viral titers at 72 h.p.i. compared to parental MVA, indicating that cloning of proteins VP2 of BTV-4 or BTV-8 weakens MVA vector replication. Co-expression of both VP2 proteins by a single rMVA did not markedly exacerbate the observed effect on MVA replication of monovalent rMVAs, although the MVA-VP2_A4_-VP2_F8_ consistently displayed lower viral titers than parental MVA at 24, 48 and 72 h.p.i. (Fig. 1C).

### Humoral immune response induced by the rMVAs

In preliminary experiments, we engineered a rMVA co-expressing proteins VP2 of BTV-1 and BTV-4 (MVA-VP2_A4_-VP2_F1_) (Supplementary Figure Fig. 1A) and evaluated its immunogenicity and protective capacity against BTV-4 in sheep. MVA-VP2_A4_-VP2_F1_ prime-boost immunization induced very low titers of nAbs against BTV-1 (PRNT_50_ ≤ 20), and no detectable nAbs against BTV-4 (Supplementary Fig. 1B). Following challenge with 10_5_ PFU of the virulent BTV-4M strain, immunized sheep did not show any degree of protection against BTV as rectal temperatures, RNAemia and hematological features were indistinguishable from those of non-immunized animals (Supplementary Fig. 1C-F). These results suggested an incompatibility between the co-expressed VP2 proteins of BTV-1 and BTV-4 in a BTV natural host. Therefore, to determine whether it was a serotype-specific incompatibility, we replaced the VP2 of BTV-1 by the VP2 protein of BTV-8.

Based on these findings, we first assessed the ability of the previously generated rMVA viral vectors to induce a BTV-nAbs in IFNAR(-/-) mice before conducting challenge experiments. Given the absence of neutralizing activity observed in sheep immunized with MVA-VP2_A4_-VP2_F1_, we also examined whether the joint administration of monovalent rMVAs (MVA-VP2_A4_ and MVA-VP2_A8_) could lead to nAbs interference. Thus, groups of IFNAR(-/-) mice (n=4) were intramuscularly immunized as indicated in the Material and Methods section, and the BTV-neutralizing activity in mice sera was evaluated.

Immunization with MVA-VP2_A4_ induced high nAbs titers against BTV-4 and the MVA-VP2_A8_ vaccine candidate elicited a strong neutralizing response against BTV-8 at 3 weeks post-boost (w.p.b.) (Fig. 2A). In contrast, mice immunized with the bivalent MVA-VP2_A4_-VP2_F8_ did not develop detectable nAbs against either serotype (Fig. 2A), consistent with supplementary data and thereby excluding its use in further challenge experiments. Importantly, no evidence of interference was observed when monovalent MVA-VP2_A4_ and MVA-VP2_A8_ were administered at separate injection sites (group MVA-VP2_A4_/MVA-VP2_A8_), which induced robust neutralizing responses against both homologous BTV serotypes (Fig. 2A). Although co-administration of monovalent rMVAs in the same inoculum (group MVA-VP2_A4_+MVA-VP2_A8_) also induced high nAb titers after the booster, PRNT_50_ values were slightly lower than those induced by the separate administration of MVA-VP2_A4_ and MVA-VP2_A8_ (Fig. 2A). Therefore, challenge experiments in IFNAR(-/-) mice and sheep were performed using separated inoculation of monovalent MVA-VP2_A4_ and MVA-VP2_A8_ during immunization.

**Figure 2.**
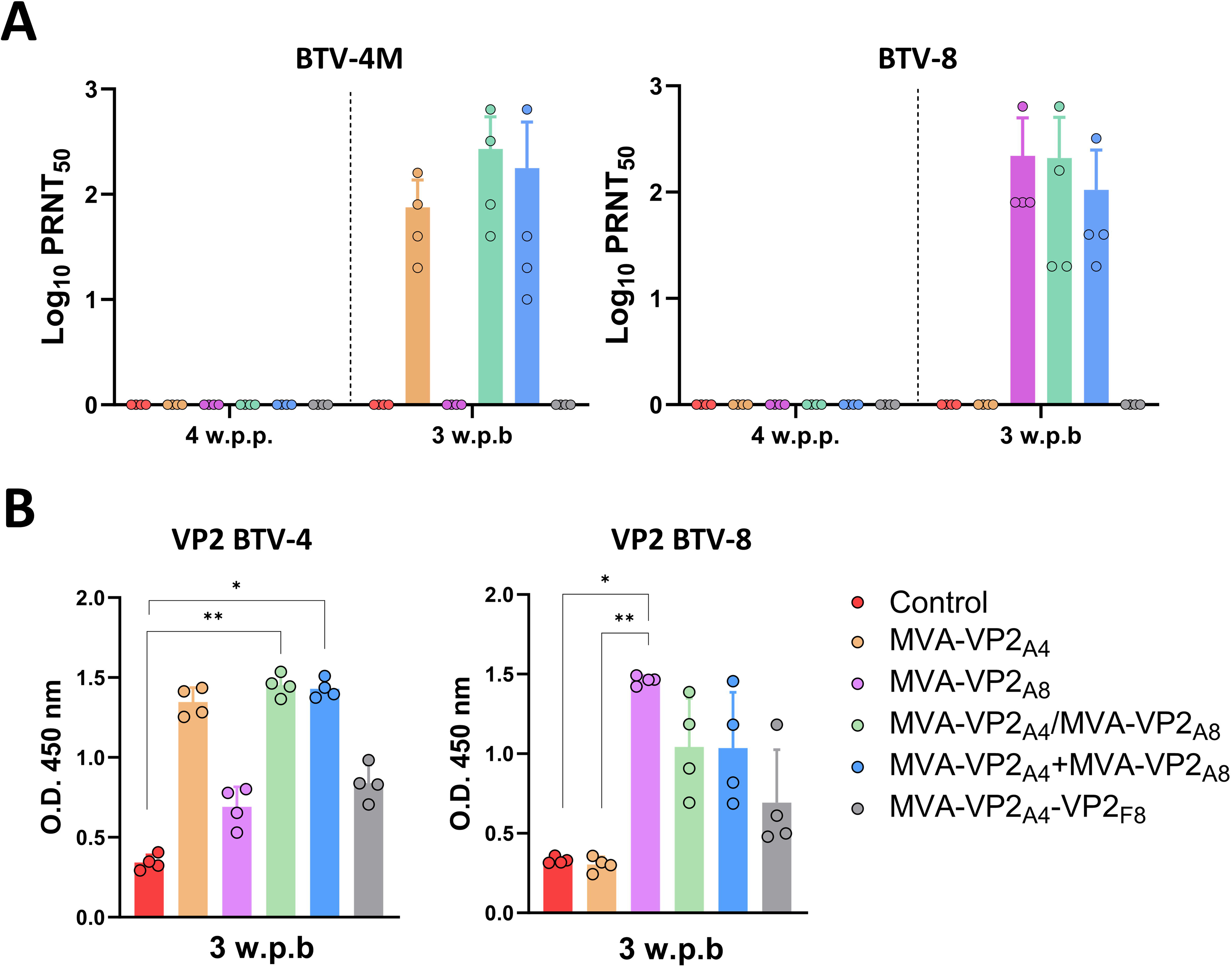
Humoral immune response induced by the rMVAs in IFNAR(-/-) mice. (A) nAbs titers against BTV-4 or BTV-8 in immunized animals by PRNT_50_ assay. Points represent individual values for each mouse, bars represent the mean values of each group and error bars represent SD. Statistical differences were found between mock-infected group and groups showing detectable nAbs titers by two-way ANOVA (post hoc Tukey test for multiple comparisons). (B) Indirect ELISA assays against VP2 of either BTV-8 or BTV-4 using sera from immunized and non-immunized animals collected at 3 w.p.b. (sera dilution 1:200). Points represent individual values for each mouse, bars represent the mean values of each group and error bars represent SD. Statistical differences were found between groups by one-way ANOVA (Kruskal-Wallis test). Group MVA-VP2_A4_/MVA-VP2_A8_ was immunized with monovalent MVA-VP2_A4_ and MVA-VP2_A8_ administered at separate injection sites.

Given that co-expression of BTV-4 and BTV-8 VP2 proteins completely abrogated the induction of BTV-neutralizing antibodies, we examined whether the MVA-VP2_A4_-VP2_F8_ was still able to elicit a non-neutralizing humoral response against both antigens. To do so, indirect ELISA and IFA assays were conducted using sera samples collected at 3 w.p.b from all immunized groups to assess VP2-specific IgG responses. Mice individually immunized with monovalent rMVA (MVA-VP2_A4_ or MVA-VP2_A8_) developed an IgG response against the homologous VP2 protein (Fig. 2B). Animals immunized with both monovalent rMVAs (groups MVA-VP2_A4_/MVA-VP2_A8_ and MVA-VP2_A4_+MVA-VP2_A8_) displayed comparable IgG levels against VP2 proteins of both serotypes (Fig. 2B). Interestingly, these two immunization groups displayed non-significantly lower levels of BTV-8 VP2-specific IgG compared to animals immunized with MVA-VP2_A8_ alone. Immunization with the bivalent MVA-VP2_A4_-VP2_F8_ also induced an IgG response specific to both VP2 proteins, although antibody levels were markedly lower than those elicited by the other immunization regimes (Fig. 2B).

Overall, our data suggest a significant interference phenomenon between heterologous VP2 proteins, being characterized by complete abolishment of BTV-neutralizing activity even with VP2 antibody recognition, which may reflect preferential focus of the humoral response to VP2 non-neutralizing epitopes.

### Protection conferred by combination of monovalent rMVAs against BTV-4M and BTV-8 in IFNAR(-/-) mice

Considering the absence of neutralizing activity after immunization with MVA-VP2_A4_-VP2_F8_, we decided to test the protective efficacy against BTV of the immunization strategy based on separated co-administration of monovalent rMVAs. To do so, four groups of IFNAR(-/-) mice (n=5) were intramuscularly immunized following a homologous prime-boost immunization regimen consisting of two doses (separate injection) of 1x10^7^ PFU of MVA-VP2_A4_ and MVA-VP2_A8,_ in a four-week interval (Fig. 3A). Three weeks post-boost, mice were subcutaneously challenged with a lethal dose of BTV-4M (10 PFU) or BTV-8 (100 PFU). Subsequently, mice were monitored for survival and RNAemia was analyzed.

**Figure 3.**
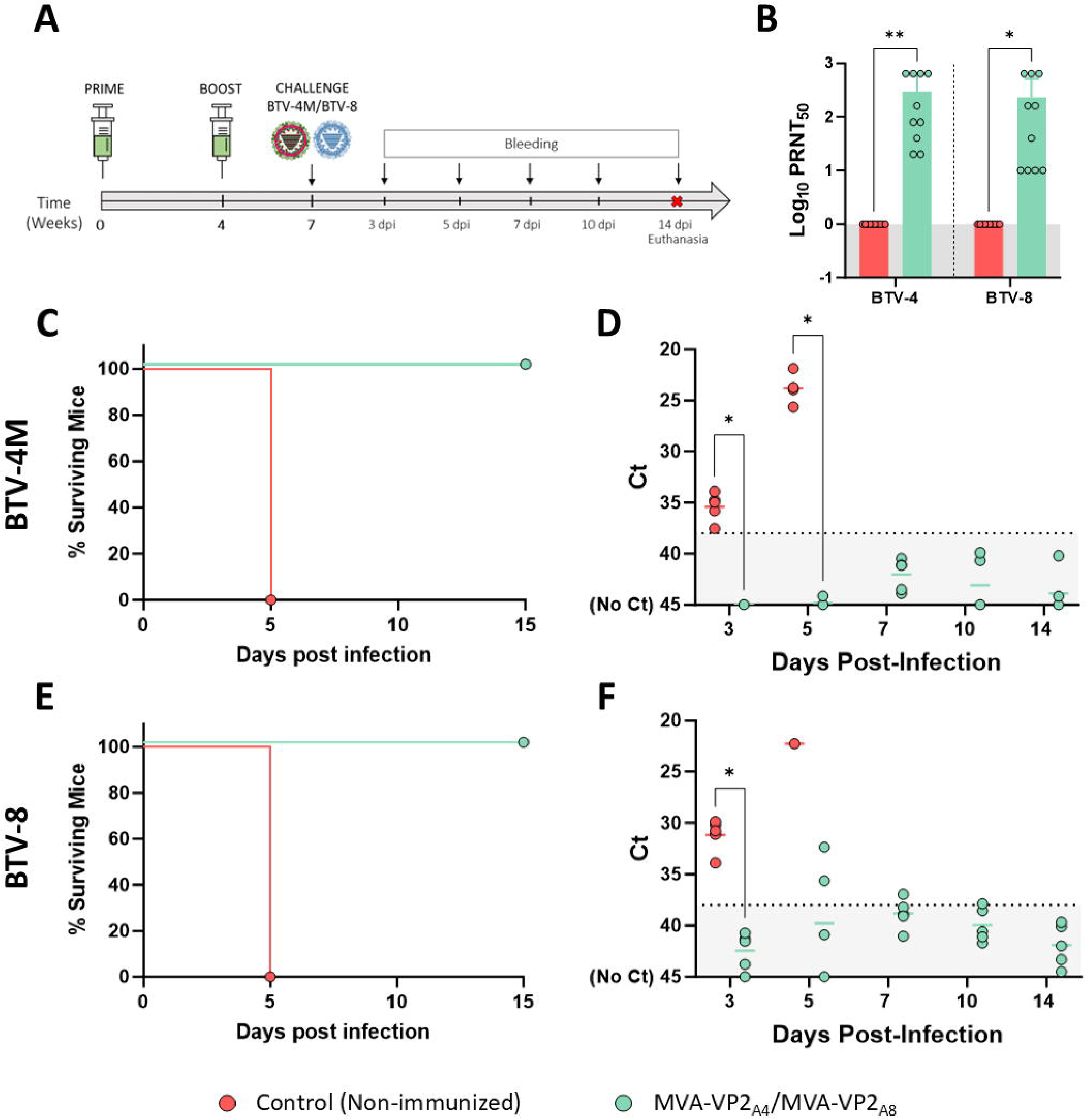
Protection of immunized IFNAR(-/-) mice against a lethal challenge with BTV-4M or BTV-8. (A) Two groups of IFNAR(-/-) mice (n=5) were immunized following a homologous prime-boost regimen consisting of two doses of MVA-VP2_A4_ and MVA-VP2_A8_ (inoculated separately). Two groups were untreated (Control). Immunized and non-immunized mice were challenged with a lethal dose of BTV-4M or BTV-8. (B) nAbs titers against BTV-4 or BTV-8 in immunized animals by plaque reduction neutralization assay. Points represent individual values for each mouse, bars represent the mean values of each group and error bars represent SD. Statistical differences between groups were analyzed using two-way ANOVA (post hoc Tukey test for multiple comparisons). (C,E) Survival rates after BTV-4M (C) or BTV-8 (E) infection. Curves were found statistically significant compared to survival curve of non-immunized mice as calculated by Log-rank test (p-value < 0.05). (D,F) RNAemia analyzed by RT-qPCR of non-immunized and immunized IFNAR(-/-) mice after viral challenge with BTV-4M (D) or BTV-8 (F). Expression of mRNA of segment 5 (encoding NS1 protein) was quantified at 3, 5, 7, 10 and 14 d.p.i. Results were expressed as Ct (left y-axis). The real-time RT-qPCR specific for BTV segment 5 was performed as described by Toussaint et al. _81_. Cut-off Ct ≥ 38 (dotted grey line). Points represent individual Ct for each mouse and lines of the corresponding color represent the mean Ct value of each group. Differences between groups were calculated by multiple t test analysis using the Sidak–Bonferroni method. * p-value < 0.05.

We measured nAbs titers in blood of immunized and non-immunized animals after the booster dose. Whereas non-immunized animals did not show any evidence of a BTV-neutralizing humoral response, mice immunized with MVA-VP2_A4_ and MVA-VP2_A8_ developed high titers of nAbs against BTV-4 and BTV-8 (Fig. 3B).

Following lethal challenge with BTV-4M, all control mice succumbed to BTV infection by day 5 post-inoculation (Fig. 3C). Furthermore, these animals displayed high levels of RNAemia on day 3 and 5 post-infection (Fig. 3D). Conversely, combined immunization with MVA-VP2_A4_ and MVA-VP2_A8_ conferred protection in immunized animals, preventing death and blocking BTV-4M replication, as no detectable RNAemia was observed at any day post-challenge (Fig. 3D).

After lethal challenge with BTV-8, non-immunized animals showed early and severe clinical signs of disease, succumbing to infection by 5 d.p.i. (Fig. 3E), and displaying RNAemia on day 3 and 5 post-infection (Fig. 3F). In contrast, 100% of immunized animals survived BTV-8 challenge (Fig. 3E). We also observed a blockade of BTV-8 replication in three out of five immunized IFNAR(-/-) mice as indicated by the absence of detectable RNA levels (Ct value ≥ 38) after challenge (Fig. 3F). Nonetheless, two out of five immunized mice exhibited detectable RNAemia at 5 d.p.i although the RNA levels were much lower than those detected in non-immunized infected mice. Anyway, these data reflect that the combination of monovalent rMVAs expressing proteins VP2 of BTV-4 and BTV-8 efficaciously protects against infections with both serotypes.

### Protective capacity of rMVAs in sheep against BTV-4M

Considering the promising results observed in IFNAR(-/-) mice, we evaluated the protective efficacy of this vaccination strategy in sheep, one of the most affected BTV natural hosts. First, we assessed the protection efficacy against BTV-4M. To do so, we immunized sheep following a prime-boost strategy consisting of two doses of MVA-VP2_A4_ and MVA-VP2_A8_ (separate injection, 1x10^8^ PFU each rMVA), administered in a four-week interval. Three weeks after the booster dose, sheep were subcutaneously challenged with 10_5_ PFU of the virulent BTV-4M strain. Thereafter, rectal temperatures, RNAemia, viraemia, hematological parameters and cytokine expression were measured at different days post-infection (Fig. 4A).

**Figure 4.**
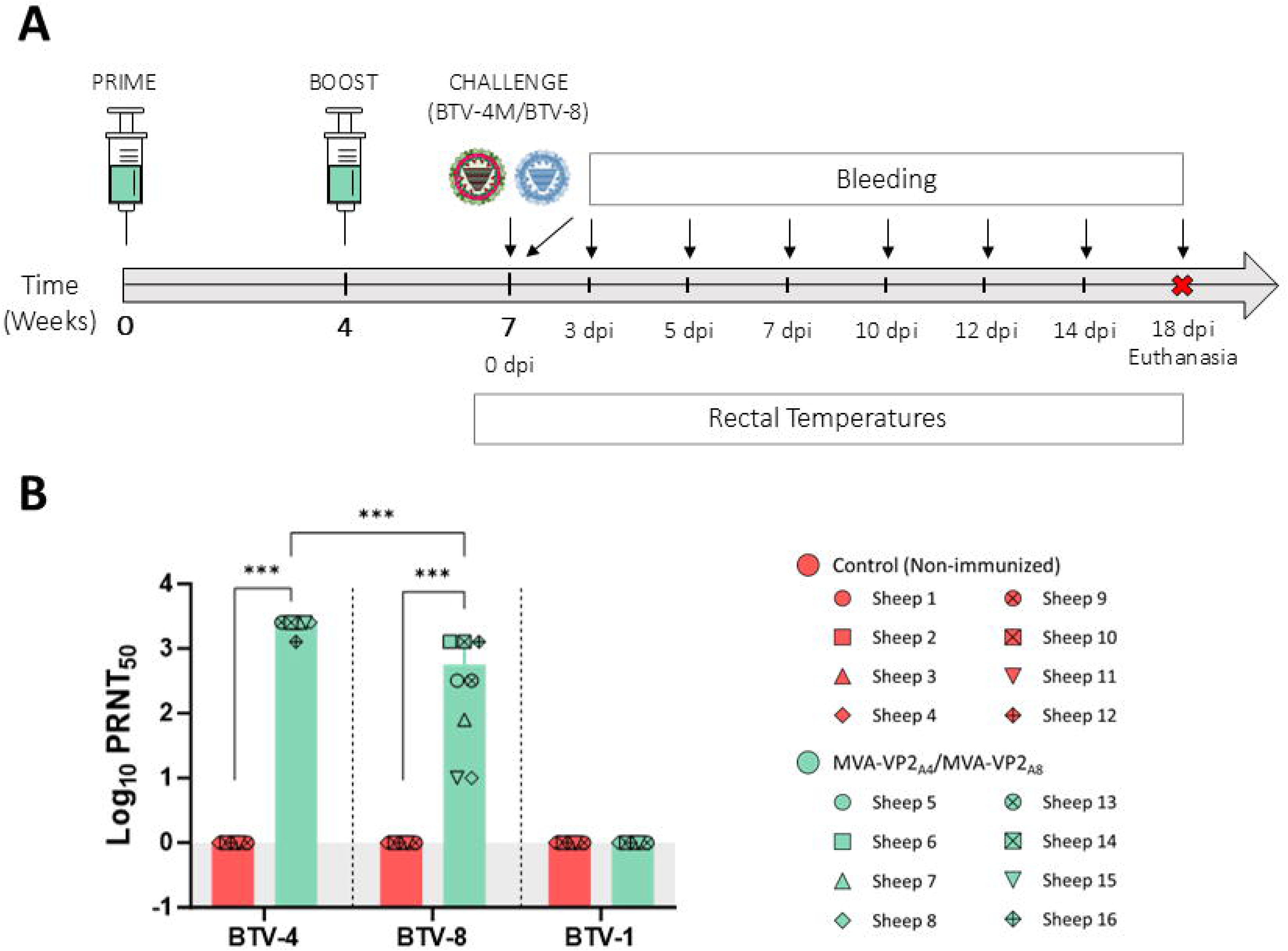
Humoral immune response in immunized sheep. (A) Groups of sheep (n=4) were immunized following a homologous prime-boost regimen consisting of two doses of MVA-VP2_A4_ and MVA-VP2_A8_ (inoculated separately). Two groups were left untreated (Control). Immunized and non-immunized sheep were challenged with BTV-4M or BTV-8. (B) nAbs titers against BTV-4, BTV-8 and BTV-1 in immunized and non-immunized sheep by PRNT_50_ assay. nAbs titers were measured in sera collected at 3 w.p.b. Bars represent the mean values of each group, points indicate the individual value of each sheep and error bars represent SD. Statistical differences were calculated by two-way ANOVA (post hoc Tukey test for multiple comparisons). *** P value < 0.001.

After the booster dose, all immunized sheep developed a strong neutralizing immune response against the homologous serotypes BTV-4 (mean PRNT_50_ titer = 1:4800) and BTV-8 (mean PRNT_50_ titer = 1:1142) (Fig. 4B). Interestingly, the neutralizing activity against BTV-8 was statistically lower (p-value < 0.0001) compared to PRNT_50_ values induced against BTV-4 (Fig. 4B). No neutralization could be detected against the heterologous serotype BTV-1 in any of the immunized sheep (Fig. 3B).

After challenge with the virulent BTV-4M, non-immunized control sheep developed increased rectal temperatures between days 6 and 8 post-infection, with some of them displaying temperature values exceeding the fever threshold established for this group (Fig. 5A). Control animals also showed detectable RNAemia levels throughout the experiment, with peak Ct values between 5 and 12 d.p.i. (Fig. 5B). Moreover, infectious virus was detected at 5 and 7 d.p.i. in some non-immunized sheep (Fig. 5C). In contrast, sheep immunized with both MVA-VP2_A4_ and MVA-VP2_A8_ showed a significant degree of protection against challenge with BTV-4M. In this sense, immunized animals presented steady rectal temperatures after challenge (Fig. 5A), showing significantly lower rectal temperature values during the febrile period of the control group (Fig. 5A). Besides, all immunized animals showed undetectable (Ct ≥ 38) levels of RNAemia and no infectious virus could be recovered from their blood after BTV-4M challenge (Fig. 5C,D), indicating a complete abrogation of BTV-4M replication in immunized sheep.

**Figure 5.**
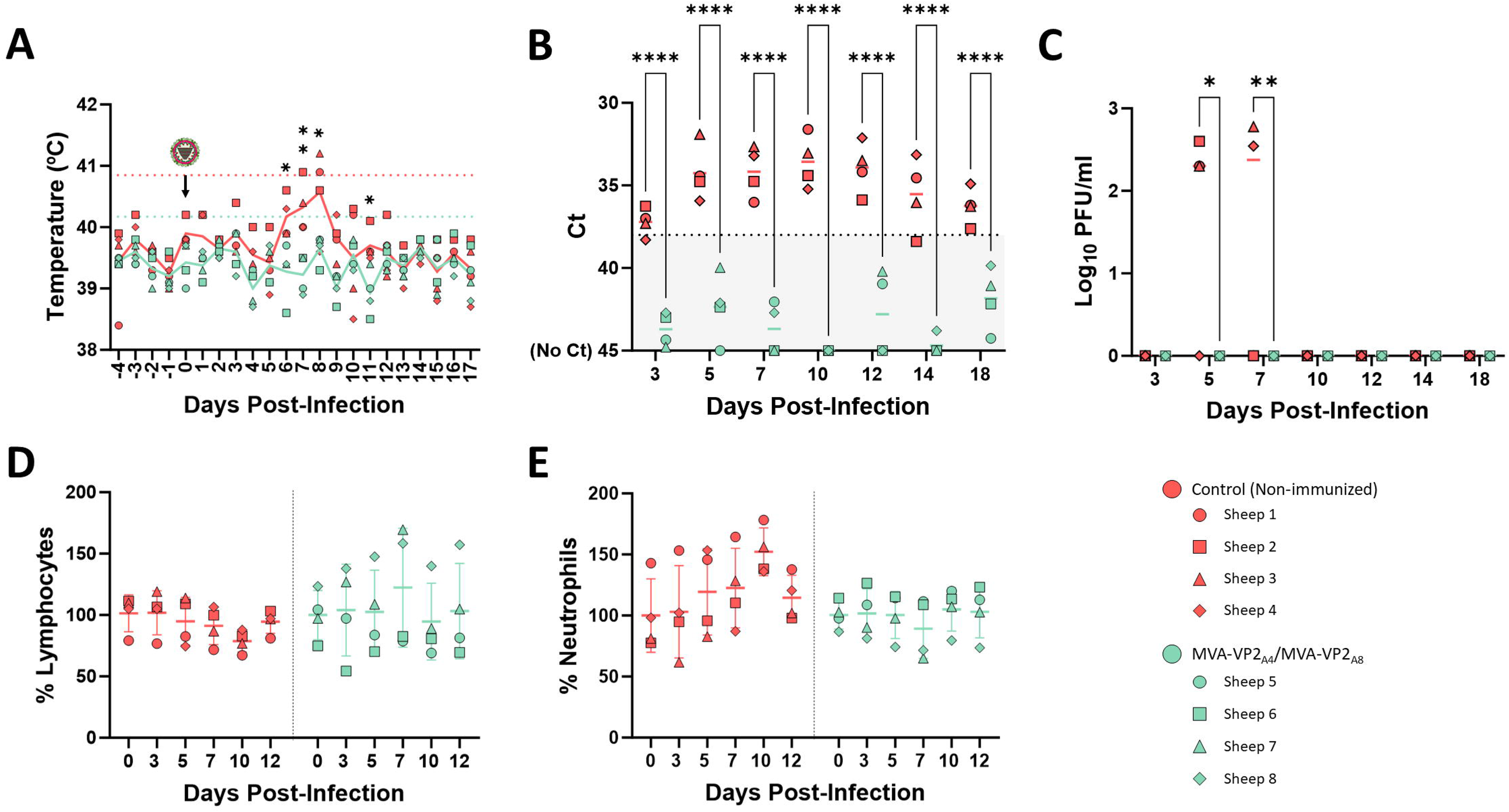
Protection of immunized sheep against a virulent challenge with BTV-4M. A group of sheep (n=4) was immunized following a homologous prime-boost regimen consisting of two doses MVA-VP2_A4_ and MVA-VP2_A8_ (inoculated separately). A group was left untreated (Control). Immunized and non-immunized sheep were challenged with BTV-4M. (A) Rectal temperatures recorded before and after BTV-4M challenge. The day of challenge (0 d.p.i.) is indicated. Points indicate the individual value of each sheep and lines represent mean rectal temperature values for each group. Coloured dotted lines represent the fever threshold for each group. (B) RNAemia analyzed by RT-qPCR of non-immunized and immunized sheep. Results were expressed as Ct (left y-axis). The real-time RT-qPCR specific for BTV segment 5 was performed as described by Toussaint et al. _81_. Cut-off Ct ≥ 38 (dotted grey line). Points represent individual Ct for each sheep and lines of the corresponding color represent the mean Ct value of each group. (C) Titers of BTV-4M recovered in sheep blood after viral challenge. Points represent individual PFU/ml value for each sheep and lines of the corresponding color represent the mean PFU/ml value of each group. (D,E) Percentages of lymphocytes and neutrophils in blood from immunized and non-immunized sheep after challenge with BTV-4M. The percentage of lymphocytes (D) and neutrophils (E) based on the total white blood cells were analyzed at days 0, 3, 5, 7, 10 and 12 post-infection. Data were normalized based on percentages recorded at 0 d.p.i. Points indicate the individual value of each sheep and lines represent the mean value of each group. * P value < 0.05, ** P value < 0.002, *** P value < 0.001, **** P value < 0.0001, using two-way ANOVA (post hoc Tukey test for multiple comparisons).

Evaluation of hematological parameters after BTV-4M challenge showed that non-immunized animals developed lymphopenia and neutrophilia between 7 and 12 d.p.i., particularly at 10 d.p.i. (Fig. 5 D,E). However, these hematologic changes were attenuated in sheep immunized with both monovalent rMVAs although one immunized animal (sheep 5) developed mild lymphopenia and neutrophilia after challenge (Fig. 5D,E).

Immunized and non-immunized sheep showed different inflammatory cytokine profiles following BTV-4M infection. Non-immunized animals exhibited higher circulating levels of IL-1α, IL-10 and IL-17 at 3 d.p.i. than those detected in immunized sheep (Fig. 6). In contrast, immunized sheep displayed higher circulating levels of IL-1β and IL-6 (Fig. 6). MVA-VP2_A4_/MVA-VP2_A8_ immunized sheep also exhibited nonsignificant lower levels of IL-1α, TNF and IL-10 at day 5 post-infection (Fig. 6). Overall, co-administration of monovalent rMVAs limited the pro-inflammatory cytokine profile associated with BTV infection.

**Figure 6.**
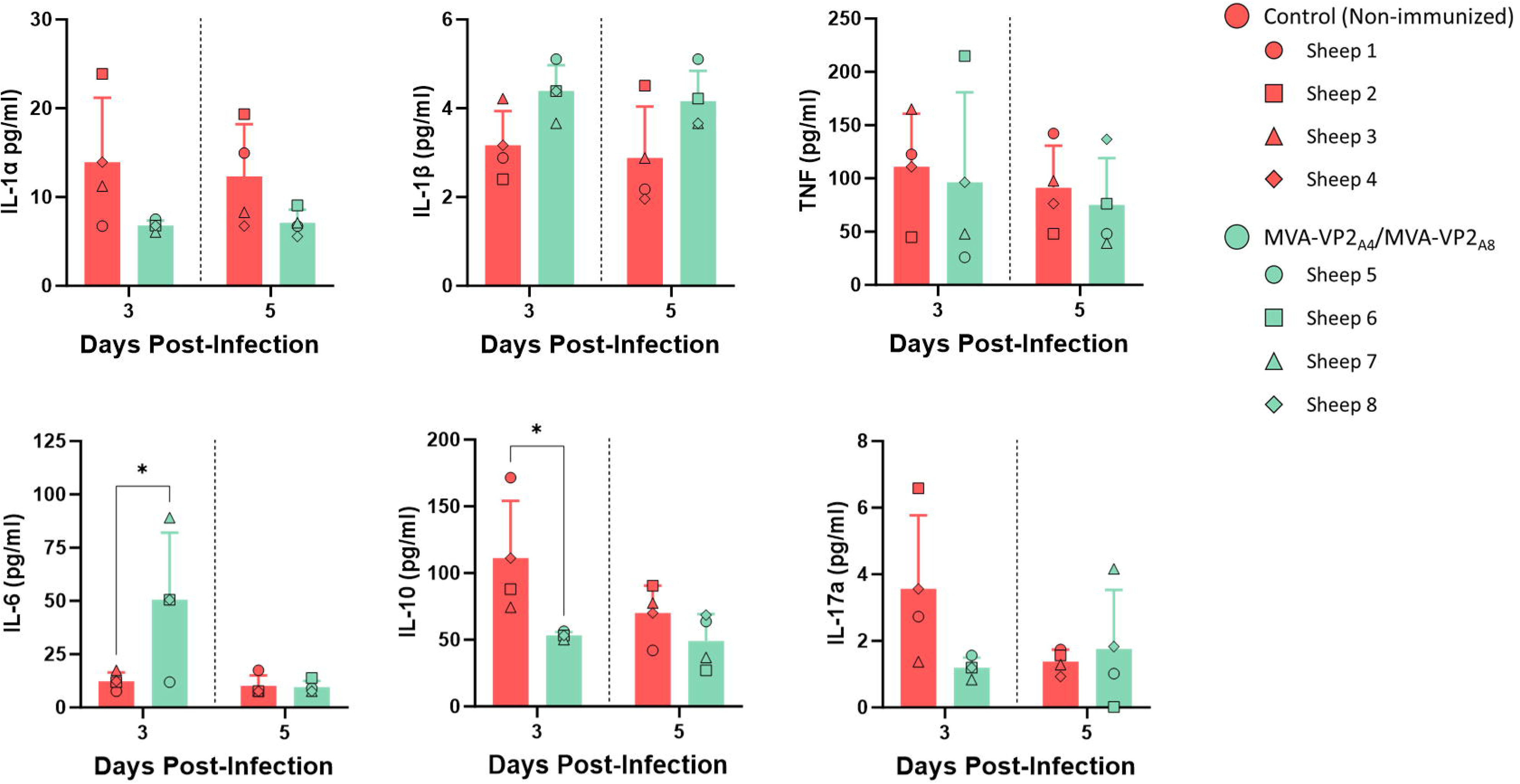
Levels of cytokines in serum samples from non-immunized and immunized sheep after challenge with BTV-4M. Sheep sera was collected at 3 and 5 d.p.i. and analyzed in Luminex immunoassays. Points represent individual values for each sheep, bars represent mean values of each group and error bars represent SD. Asterisks denote significant differences between immunized and non-immunized sheep. * P value < 0.05, using two-way ANOVA (post hoc Tukey test for multiple comparisons).

### Protective capacity of rMVAs in sheep against BTV-8

Thereafter, we evaluated the capacity of this vaccination strategy to elicit protection against BTV-8 in sheep. To do so, we followed the same immunization regime and challenged immunized and non-immunized sheep with a subcutaneous dose of 10_5_ PFU of BTV-8. Afterwards, rectal temperatures, RNAemia, viraemia, hematological parameters and cytokine expression were measured at different days post-infection (Fig. 4A).

Similarly to BTV-4M infection, non-immunized sheep showed a marked increase in their rectal temperatures between days 6 and 8 post-infection, with some of them displaying temperatures above the fever threshold (Fig. 7A). On the contrary, immunized sheep did not exhibit abnormal rectal temperature increases at any time point evaluated, showing stable temperatures throughout the experiment, except for immunized sheep 16 that reached 40.8°C (above the fever threshold) at 6 d.p.i. (Fig. 7A). Non-immunized animals also displayed detectable levels of RNAemia from day 5 post-infection, peaking between 7 and 12 d.p.i. (Fig. 7B), which coincides with abnormally increased rectal temperatures. Moreover, we detected high titers of infectious virus in blood of control animal at 5, 7 and 10 d.p.i. (Fig. 7C). Despite combined immunization with MVA-VP2_A4_ and MVA-VP2_A8_ significantly curbed the high degree of viral replication observed in control animals after BTV-8 challenge, all four immunized sheep showed detectable RNAemia at day 5 post-infection (Fig. 7B). Nonetheless, these animals were aRNAemic (Ct value ≥ 38) in subsequent days until the end of the experiment and no infectious virus was detected in their blood after BTV-8 challenge (Fig. 7C). Overall, these data also highlight the capacity of the vaccination strategy to prevent viral replication after BTV-8 challenge.

**Figure 7.**
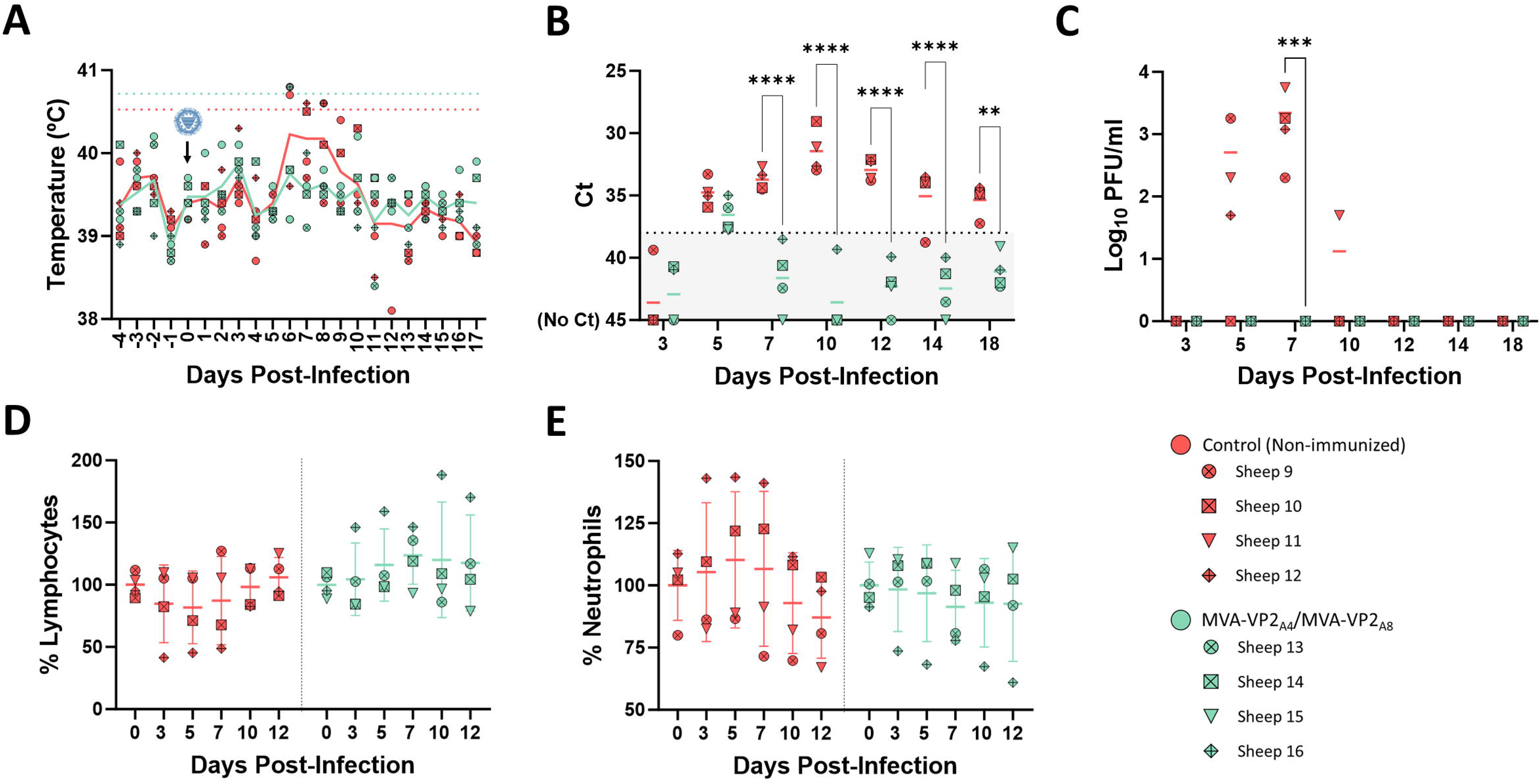
Protection of immunized sheep against a virulent challenge with BTV-8. A group of sheep (n=4) was immunized following a homologous prime-boost regimen consisting of two doses of MVA-VP2_A4_ and MVA-VP2_A8_ (inoculated separately). A group was left untreated (Control). Immunized and non-immunized sheep were challenged with BTV-8. (A) Rectal temperatures recorded before and after BTV-8 challenge. The day of challenge (0 d.p.i.) is indicated. Points indicate the individual value of each sheep and lines represent mean rectal temperature values for each group. Coloured dotted lines represent the fever threshold for each group. (B) RNAemia analyzed by RT-qPCR of non-immunized and immunized sheep. Results were expressed as Ct (left y-axis). The real-time RT-qPCR specific for BTV segment 5 was performed as described by Toussaint et al. _81_. Cut-off Ct ≥ 38 (dotted grey line). Points represent individual Ct for each sheep and lines of the corresponding color represent the mean Ct value of each group. (C) Titers of BTV-8 recovered in blood of sheep after viral inoculation. Points represent individual PFU/ml value for each sheep and lines of the corresponding color represent the mean PFU/ml value of each group. (D, E) Percentages of lymphocytes and neutrophils in blood from immunized sheep after challenge with BTV-8. The percentage of lymphocytes (D) and neutrophils (E) based on the total white blood cells were analyzed at days 0, 3, 5, 7, 10 and 12 post-infection. Data were normalized based on percentages recorded at 0 d.p.i. Points indicate the individual value of each sheep and lines represent the mean value of each group. ** P value < 0.002, *** P value < 0.001, **** P value < 0.0001, using two-way ANOVA (post hoc Tukey test for multiple comparisons).

Analysis of hematological parameters revealed heterogenic results for both immunized and non-immunized sheep groups after challenge with BTV-8. Whereas control sheep 9 and 11 showed steady hematologic values, control sheep 10 and 12 suffered a strong reduction in the percentage of lymphocytes as well as a marked increase in the percentage of neutrophils after BTV-8 challenge (Fig. 7D,E). In contrast, immunized sheep did not suffer from lymphopenia or neutrophilia as they displayed steady values after challenge, except for immunized sheep 14 that slightly suffered these hematologic features at day 3 and 5 post-infection (Fig. 7D,E).

Analysis of circulating cytokines revealed similar circulating cytokine profiles between groups after BTV-8 infection. Immunized sheep displayed lower levels of IL-1α at 3 d.p.i. and increased levels of circulating IL-6 at 3 and 5 d.p.i. compared to the control group (Fig. 8). Overall, we did not observe significant changes regarding levels of circulant IL-1α, IL-1β, TNF, IL-6, IL-10 and IL-17 between the immunized and control group at 3 or 5 d.p.i. (Fig. 8), which is consistent with similar viral replication levels between groups at the beginning of infection.

**Figure 8.**
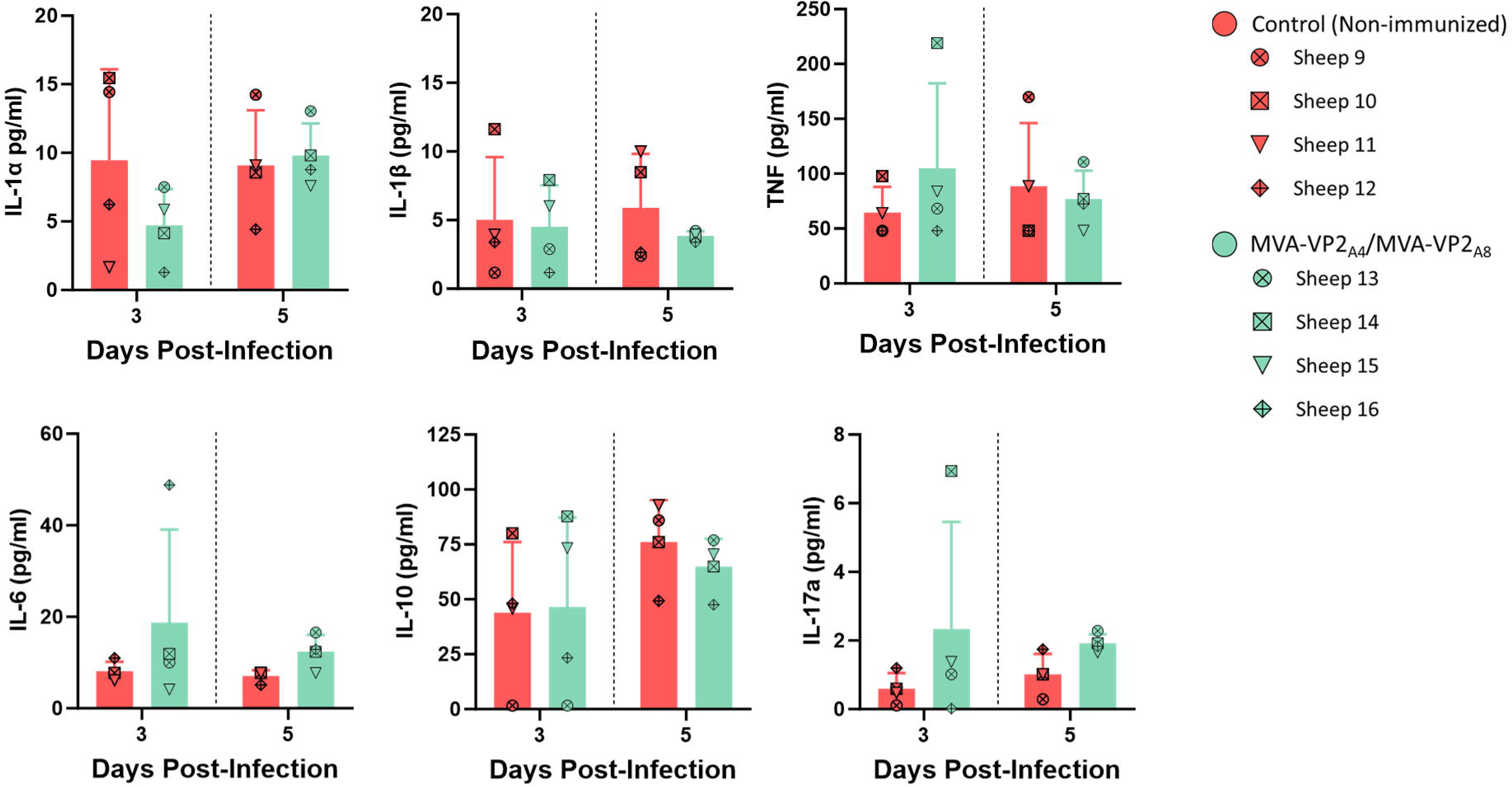
Levels of cytokines in serum samples from non-immunized and immunized sheep after challenge with BTV-8. Sheep sera were collected at 3 and 5 d.p.i. and analyzed in Luminex immunoassays. Points represent individual values for each sheep, bars represent mean values of each group and error bars represent SD. Asterisks denote significant differences between immunized and non-immunized sheep. * P value < 0.05, using two-way ANOVA (post hoc Tukey test for multiple comparisons).

## DISCUSSION

Historically, classical live-attenuated and inactivated vaccines have been the cornerstone of BTV control. However, these conventional approaches frequently lack DIVA character, involve safety concerns due to potential reversion to virulence or genomic reassortment, and fail to provide immunity against the plethora of BTV serotypes ^50,82^. Recent advancements in innovative vaccination approaches aim to manage this economically significant disease by providing next-generation vaccines that address safety, efficacy, and DIVA characteristics ^50–52,65,83^. Diverse experimental vaccine candidates demonstrate promising improvements compared to LAVs or inactivated vaccines, particularly the induction of broad protective immunity against multiple BTV serotypes, but none have been licensed yet ^59,60,84^. LAVs and inactivated BTV vaccines base their protective efficacy on the induction of a potent humoral immune response directed against protein VP2 ^50^, possessing conformational nAbs-inducing epitopes within its trimeric configuration ^85–87^. Most new-generation BTV vaccines also base their protective efficacy on the formulation of protein VP2 ^50–52,88^, highlighting the importance of this antigen to provide protective immunity. Consequently, inclusion of VP2 remains essential for the rational design of newly generated BTV vaccines. Here, we explored a novel MVA-based vaccination approach that aimed to provide robust bivalent protection against BTV based on the co-expression of proteins VP2 from different BTV serotypes.

The MVA viral vector allows simultaneous expression of different heterologous genes, which is ideal to formulate different antigens of one pathogen or to design multipathogen vaccines by using one single viral vector ^58,89^. Thus, our main goal was the design of a single rMVA able to co-express two proteins VP2 from different BTV serotypes, an approach that would be useful for dealing with epizootic BTV events that involve more than one serotype. Surprisingly, evaluation of bivalent VP2-based rMVAs as vaccine vectors showed a significant interference between VP2 proteins from different BTV serotypes that critically affected their reliability as protective orbiviral antigens. An impaired BTV-neutralizing activity characterized the immune response induced by the MVA-VP2_A4_-VP2_F8_ but also the immunogenicity of another bivalent rMVAs, such as MVA-VP2_A4_-VP2_F1_, or another rMVA that expressed proteins VP2 of BTV-4 and BTV-8 but cloned in different loci (*F13L* and *TK* loci) (data not shown). Although this interference should be evaluated using another expression platform, the concordance observed between different rMVAs suggests that the absence of neutralizing activity is not due to aspects related to the MVA viral vector, even more considering that non-neutralizing humoral immune responses against proteins VP2 were observed, confirming antigen expression. Our results indicate that antigen expression occurs, but conformational neutralizing epitope integrity could be compromised, which may suggest potential misfolding or degradation of VP2 proteins within MVA-infected cells. Heat-shock protein 90 (HSP90) prevents BTV protein degradation within infected cells ^90^. Competition for chaperons between proteins VP2 could theoretically lead to misfolding, formation of mixed or unstable oligomers, and subsequent targeting of proteins VP2 to degradation pathways. This would not impede antigen presentation and induction of non-neutralizing antibodies, but it would disrupt conformational neutralization determinants within protein VP2, which is consistent with the observed non-neutralizing antibody responses against both VP2 proteins. Alternatively, structure-guided engineering of protein VP2 may help inhibit the observed interference. Rational design of orbiviral protein VP2 enhanced protein stability, improved protein folding and preserved neutralizing epitope architecture ^91^. Applying a similar approach to BTV VP2 protein may reduce misfolding or aberrant oligomerization, maintaining key conformational determinants even under simultaneous expression of VP2 proteins.

The interference phenomenon observed between VP2 antigens may be related to structural features intrinsic to VP2. This outer-capsid protein forms triskelion-shaped trimers that are essential for virus assembly, receptor recognition, and infectivity ^28^. This trimeric architecture organization not only contributes to receptor binding and host cell attachment but also influences the spatial arrangement, accessibility, and stability of neutralizing epitopes located mainly within the flexible Tip domain, thereby affecting immune recognition _30,_^92^. The central hub domain of each VP2 monomer mediates the trimerization, and constitutes the main intersubunit contact region ^28^. Structural predictions indicated that the VP2 hub domains from BTV-4 and BTV-8 retain a highly conserved fold (Fig. 9A), suggesting that heterologous interactions between monomers are structurally feasible. Therefore, co-expression of VP2 proteins from different serotypes in the same cell and in the absence of the remaining viral structural proteins, could allow the formation of heterologous VP2 trimers. Such heterologous interactions could disrupt the three-fold symmetry found in homologous trimers, potentially altering the relative positioning of the flexible Tip domain and modifying the presentation of neutralizing epitopes. Thus, the aberrant disposition of the Tip domain in these heterologous trimers would affect the neutralizing immune response elicited against these antigens. Antibodies elicited under these conditions might preferentially recognize conserved or structurally preserved regions of VP2 while exhibiting reduced neutralizing activity due to altered exposure of serotype-specific epitopes. However, the interface analyses performed using PDBePISA provide important context for interpreting these structural predictions. Although homologous and heterologous VP2 assemblies displayed conserved electrostatic interaction networks involving complementary Lys/Arg and Asp/Glu residues (Fig. 9B-E, Table S1), none of the predicted trimers exhibited physicochemical characteristics typically associated with stable biological oligomers. All interfaces were characterized by small buried surface areas (approximately 850–1070 Å²), a limited number of interface residues (approximately 22–32 per chain), modest networks of hydrogen bonds and salt bridges (approximately 12–15 interactions per interface), low solvation energy gains (−2 to −4 kcal/mol), and Complex Formation Significance Scores (CSS) of 0 (Table S2). Together, these parameters indicate that both homologous and heterologous contacts represent weak interactions that are unlikely to support stable VP2 trimerization as independent assemblies. These findings suggest an important distinction between structural compatibility and biological relevance. While electrostatic complementarity is sufficient to permit intermolecular contacts between VP2 monomers from different serotypes, such interactions alone do not generate the stability required for functional oligomerization. Instead, VP2 trimer stability within the virion is likely achieved through a highly cooperative assembly process involving additional structural components. In the native BTV particle, VP2 is constrained by the geometry of the icosahedral capsid and forms extensive contacts with VP5 and the underlying VP7/VP3 scaffold, creating a multivalent interaction network that provides both geometric precision and energetic stabilization. Consistent with this interpretation, experimental studies ^30^ have shown that VP2 expressed in isolation is predominantly detected as monomers and other weakly associated oligomeric forms, whereas trimeric species represent only a fraction of the protein population. Considering that protein VP5 stabilizes VP2 trimers through its interaction with the central hub domain ^28^, inclusion of VP5 in the vaccine formulation may promote more native-like VP2 organization and improve neutralizing antibody induction. Such an approach is complicated by the predominantly serotype-specific nature of VP2–VP5 interactions ^84,93–95^, and would require the incorporation of additional heterologous genes into the vaccine platform. However, this VP2-VP5 specificity could promote homologous trimers arrangement in the context of different VP2 proteins co-expression. In summary, our structural prediction analysis indicates that although heterologous VP2 monomers can establish chemically compatible contacts, these interactions remain weak and biologically insignificant in the absence of the native capsid environment. In any case, VP2 expression in expression alone would lead to a minority trimeric form and a predominancy of monomeric configurations along with other weakly associated configurations, that are able to retain the nAbs induction capacity ^30^. Collectively, these observations argue against structural interference through stable heterologous trimer formation as the primary explanation for the complete loss of neutralizing activity observed during immunization. Hence, the observed interference is more likely to arise from immunological mechanisms affecting antigen processing, competition, or immune recognition than from the formation of stable mixed VP2 assemblies.

**Figure 9.**
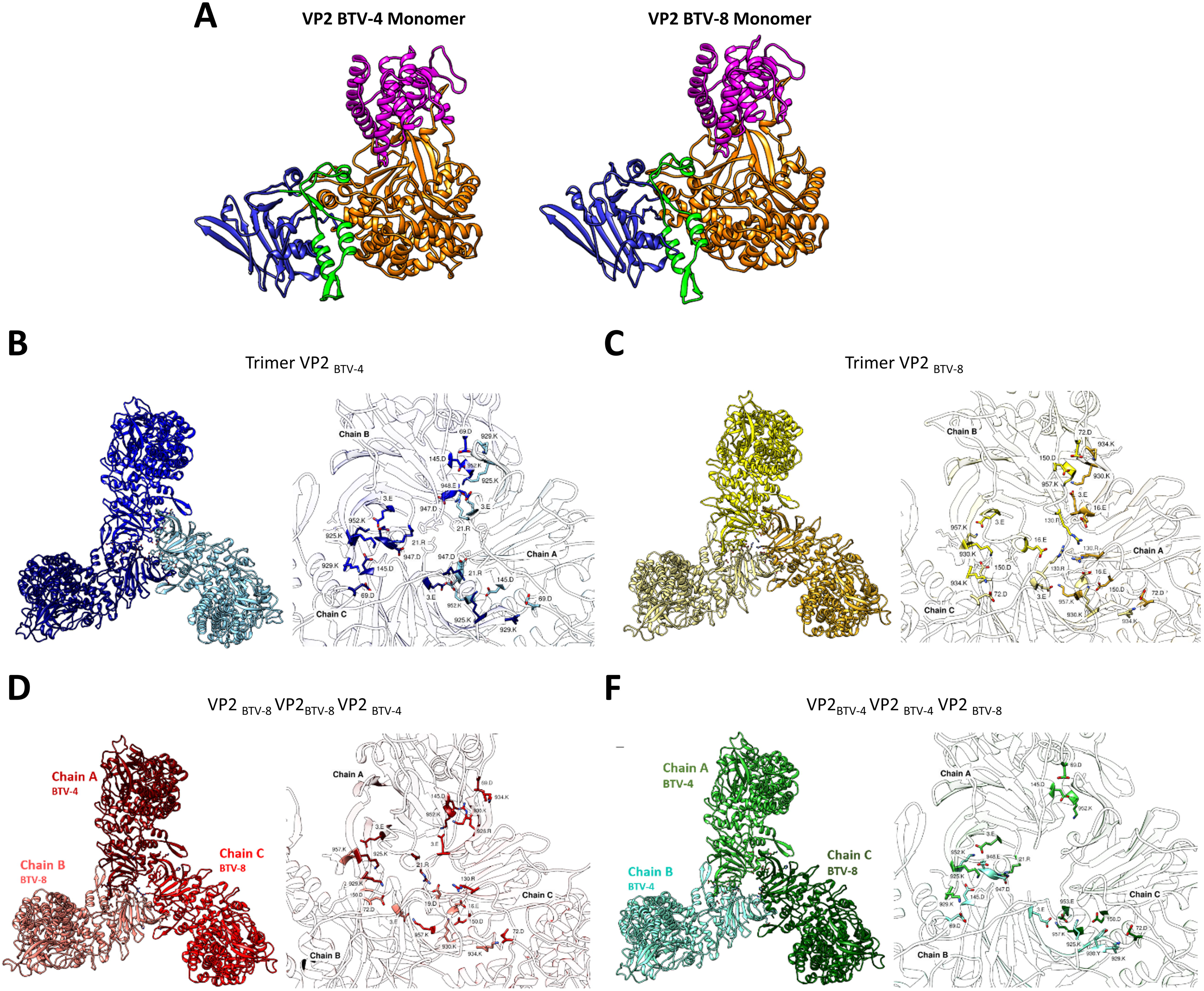
Structure prediction of the receptor-binding protein VP2 of BTV-4 and BTV-8. (A) Top view ribbon model of the VP2 monomer colored by its four domains: hub in blue, hairpin (50–120) in green, body (163–190, 408–849) in orange and external tip body (191-407) in magenta. Right panel represents the VP2 BTV-4 monomer and the left panel the VP2 BTV-8 monomer. (B-E) Ribbon model, viewed from top view, of the hub domain interfaces for each structure prediction: BTV-4 homotrimer (B), BTV-8 homotrimer (C), heterologous trimer (1xBTV-4, 2xBTV-8) (D) and heterologous trimer (2xBTV-4, 1xBTV-8) (E). Principal salt bridges contacts identified using PDBePISA analysis are highlighted.

Sequential exposure to highly related viral antigens can bias the humoral response toward conserved, cross-reactive epitopes that are often poorly neutralizing, which limits serotype- or strain-specific neutralizing antibody responses as documented for dengue virus, influenza virus or SARS-CoV-2 virus ^96–102^. Although immunological imprinting due to repeated exposure to antigenically related proteins is a well-known phenomenon, the effect on the antibody response of simultaneous immunization with multiple immunogens is not well-known. Simultaneous processing of multiple antigens can lead to changes on the response against their epitopes, with distortion on epitope selection and immunodominance ^103,104^. Therefore, antigenic interference may arise when related antigens are presented simultaneously. Thus, an alternative explanation for the critical loss of VP2-induced neutralizing responses could be related with preferential recognition of shared regions between VP2 proteins. If it were the case, rational engineering of protein VP2 to remove conserved, non-relevant regions could help redirect the response toward relevant neutralizing epitopes.

Alternative to bivalent VP2-expressing rMVAs, we explored combination of monovalent rMVA expressing proteins VP2 of different serotypes. Co-administration of monovalent rMVAs significantly blocked homologous BTV replication, inducing robust protection in a highly susceptible BTV host such as sheep. Our strategy is comparable to field implementation of multivalent cocktails of conventional vaccines _105_. Similarly, next-generation vaccine approaches such as VLPs and engineered-LAV DISA/DISC vaccine platforms also rely on co-administration of monovalent constructs delivering or expressing serotype-specific VP2 proteins to achieve broad-spectrum protection against BTV ^67,69,72,84,106^. Despite being highly effective against BTV, the constraint of formulating serotype-specific particles to achieve multiserotype protection adds complexity to their implementation as vaccines against BT. Cocktail of DISA/DISC vaccines, although multiserotype protective, rely on replacing serotype-specific VP2 and VP5 proteins on a common viral backbone, a process that may lead to generation of low-fitness recombinant viruses that could limit replication efficiency, reduce antigen yield and, ultimately, constrain large-scale production. VLPs need adjuvants to achieve robust immunity ^33^ and face potential challenges related to protein complex instability as they require correct assembly of proteins VP2, VP5, VP7 and VP3, which might be a bottleneck for production and field application. Co-administration of monovalent rMVA-based vaccines circumvents these issues by avoiding particle instability or full-virus engineering as the highly stable MVA vaccine platform enables straightforward multiserotype vaccine combinations and benefit from highly standardized, scalable manufacturing pipelines already established for other rMVA vaccines ^107–110^, ensuring consistent antigen expression and robust batch-to-batch reproducibility. Therefore, our vaccination strategy can be considered a robust DIVA-compatible alternative to available LAV or inactivated vaccines that can be tailored to specific multiserotype BTV epizootics.

Although significant protection is achieved after immunization with both monovalent rMVAs, our data show an imbalanced response in sheep towards protein VP2 of BTV-4, resulting in higher nAbs titers against this serotype and complete protection in absence of viral replication after challenge. In contrast, the response induced by the MVA-VP2_A8_ induced lower nAbs titers and viral replication occurred early after challenge, suggesting a biased humoral response towards neutralizing epitopes of VP2 protein of BTV-4. Without ignoring a possible methodological error during sheep immunization or the potential impact of BTV-8 VP2 protein expression on MVA fitness that may lead to a lessened exposure of this antigen to the immune system, the imbalanced neutralizing response directed against one of these antigens might reflect immunological dominance of BTV-4 VP2 over BTV-8 VP2. A similar pattern of serotype-specific VP2-biased immunodominance/interference has been reported in other BTV multivalent vaccine strategies. Pérez de Diego, et al. (2011) suggested antigenic interference between VP2 proteins after immunization of sheep with a combination of VLPs of BTV-1 and BTV-4 _69_. Likewise, bivalent immunization of cattle with BTV-2 and BTV-4 inactivated vaccines led to higher nAbs titers against BTV-2 ^111^. Altogether, these findings indicate that protein VP2 of certain serotypes may act as immunodominant antigens, which may be related to structural or antigenic features that confer immunodominance when co-delivered with other VP2 antigens of distinct serotypes. In any case, it illustrates another factor to consider when formulating multivalent VP2-based vaccines against orbiviruses.

The predominant serotype-specificity of the immune response directed against protein VP2 remains as a major constrain for the development of broad-spectrum vaccines against BTV ^35,112^. Beyond this well-known limitation, VP2-based vaccine development must face additional challenges such as potential antibody-dependent enhancement of infection (ADE) phenomena ^113^, or the emergence of immunodominance between VP2 antigens ^69,111^. The unfeasibility of designing vaccine platforms that co-express a variety of proteins VP2 adds a significant factor to be considered regarding the formulation of this protein as a vaccine antigen. Despite further research is required to fully elucidate the mechanism underlying the described vaccine interference, our findings reflect critical considerations for future development of next-generation vaccines against BTV (and related orbiviruses) and represent a novel example of antigenic interference in the context of global vaccine development.

### AUTHOR CONTRIBUTIONS

J.O. and L.J.C. conceived the study. L.J.C. wrote the manuscript with contributions from D.A.C., J.S.P., J.O., and R.B. J.S.P., M.M.L. and R.B. generated the rMVAs and L.J.C. characterized them. L.J.C., I.M.A., S.U.T. and E.C.P. conducted mice experiments. J.B.S. performed sheep immunization. L.J.C., I.M.A., A.N., J.B.S. and E.C.P. analyzed antibodies, RNAemia, viremia and hematological parameters in mice and sheep. E.C.P. conducted cytokine analysis. D.A.C. performed bioinformatic analysis.

## ACKNOWLEDGEMENTS

We thank the animal facility and the veterinary unit of CISA-INIA/CSIC for animal handling and technical assistance.

---

**Supplementary Figure 1.**
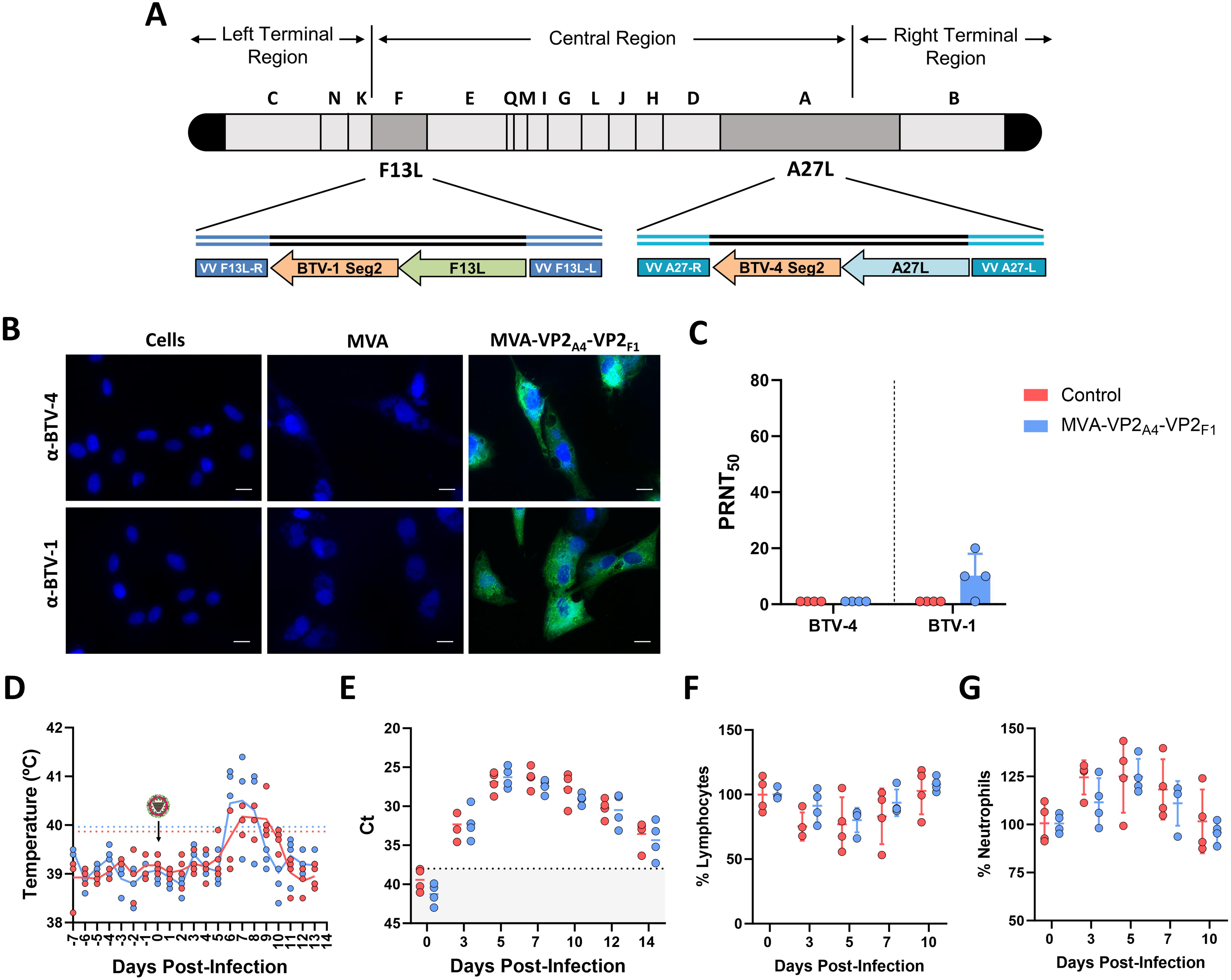
Expression analysis and protective efficacy of the MVA-VP2_A4_-VP2_F1_ in sheep. (A) Schematic representation of the cloning strategy within the MVA genome. Segment 2 of BTV-1 and BTV-4 were cloned in the F13L and A27L loci, respectively. (B) Indirect immunofluorescence of DF-1 cells infected (MOI = 0.1) with parental MVA, MVA-VP2_A4_-VP2_F1_ or non-infected (control). Proteins VP2 of BTV-4 and VP2 of BTV-1 (green) were detected using mouse polyclonal sera against BTV-4 or BTV-1. Nuclei were stained with DAPI. Scale bars 20 µm. (B,C,D,E,F) A group of sheep (n=4) was immunized following a homologous prime-boost regimen consisting of two doses of MVA-VP2_A4_-VP2_F1_. A group was left untreated (Control). Immunized and non-immunized sheep were challenged with BTV-4M. (C) nAbs titers against BTV-4, BTV-8 and BTV-1 in immunized and non-immunized sheep by PRNT_50_ assay. nAbs titers were measured in sera collected at 3 w.p.b. Points represent individual values for each sheep, bars represent the mean values of each group and error bars represent SD. (D) Rectal temperatures recorded before and after BTV-4M challenge. The day of challenge (0 d.p.i.) is indicated. Points represent individual values, lines represent mean rectal temperature values for each group and error bars represent SD. Coloured lines represent the fever threshold for each group. (E) RNAemia analyzed by RT-qPCR of non-immunized and immunized sheep. Results were expressed as Ct (left y-axis). The real-time RT-qPCR specific for BTV segment 5 was performed as described by Toussaint et al. (88). Cut-off Ct ≥ 38 (dotted grey line). Points represent the individual Ct values of each group, bars represent mean Ct values and error bars represent SD. (F,G) Percentages of lymphocytes and neutrophils in blood from immunized and non-immunized sheep after challenge with BTV-4M. The percentage of lymphocytes (F) and neutrophils (G) based on the total white blood cells were analyzed at days 0, 3, 5, 7, 10 and 12 post-infection. Data were normalized based on percentages recorded at 0 d.p.i. Points represent the individual values of each group, bars represent mean values of each group and error bars represent SD. No statistical differences were found groups by two-way ANOVA (post hoc Tukey test for multiple comparisons).

**Table S1.** Major hydrogen bonds and salt bridges identified in homologous and heterologous VP2 trimers*.

| Trimer | Interface | Major salt bridges | Representative hydrogen bonds |
|---|---|---|---|
| **BTV-4 homotrimer** | A-B | Asp69-Lys929; Asp145-Lys925; Asp947/Glu948-Arg21; Glu3-Lys952 | Tyr126-His125; Tyr126-Glu124; Asn955-Arg921; Asn955-Glu924; Asn946-Glu3 |
|  | B-C | Asp69-Lys929; Asp145-Lys925; Asp947/Glu948-Arg21; Glu3-Lys952 | Tyr126-His125; Tyr126-Glu124; Asn955-Arg921; Asn955-Glu924; Asn946-Glu3 |
|  | A-C | Asp69-Lys929; Asp145-Lys925; Asp947/Glu948-Arg21; Glu3-Lys952 | Tyr126-His125; Tyr126-Glu124; Asn955-Arg921; Asn955-Glu924; Asn946-Glu3 |
| **BTV-8 homotrimer** | A-B | Lys930-Asp150; Lys934-Asp72; Glu3-Lys957; Glu16-Arg130 | Lys929-Asn960; Ser131-Glu16; Ser146-Leu17; Gly20-Lys147 |
|  | B-C | Lys930-Asp150; Lys934-Asp72; Glu3-Lys957; Glu16-Arg130 | Lys929-Asn960; Ser131-Glu16; Ser146-Leu17; Gly20-Lys147 |
|  | A-C | Lys930-Asp150; Lys934-Asp72; Glu3-Lys957; Glu16-Arg130 | Lys929-Asn960; Ser131-Glu16; Ser146-Leu17; Gly20-Lys147 |
| **Heterologous trimer<br>(1×BTV-4 + 2×BTV-8)** | A-B (4/8) | Lys925-Asp150; Lys929-Asp72; Arg21-Asp19; Glu3-Lys957 | Tyr22-Glu953; Arg921-Asn960; Glu924-Asn960; Glu124-Arg130 |
|  | A-C (4/8) | Asp145-Lys930; Asp69-Lys934; Lys952-Glu3 | Asn946-Glu3; Asn955-Lys929; Thr141-Leu17; Glu948-Tyr21 |
|  | B-C (8/8) | Lys930-Asp150; Lys934-Asp72; Glu3-Lys957; Glu16-Arg130 | Tyr21-Glu953; Arg926-Asn960; Lys929-Asn960; Leu17-Ser146 |
| **Heterologous trimer<br>(2×BTV-4 + 1×BTV-8)** | A-B (4/4) | Asp69-Lys929; Asp145-Lys925; Asp947/Glu948-Arg21; Glu3-Lys952 | Tyr22-Glu948; Arg921-Asn955; Glu924-Asn955; Leu18-Thr141 |
|  | A-C (4/8) | Asp145-Lys930; Asp69-Lys934; Lys952-Glu3 | Asn946-Glu3; Asn955-Lys929; Thr141-Leu17; Glu948-Tyr21 |
|  | B-C (4/8) | Lys925-Asp150/Glu953; Lys929-Asp72; Glu3-Lys957 | Tyr22-Glu953; Arg921-Asn960; Glu924-Asn960; Leu18-Ser146 |

\*Hydrogen bonds and salt bridges were identified using PDBePISA. Only the most recurrent and structurally relevant contacts are shown.

**Table S2.** Global summary of VP2 interface properties in homologous and heterologous trimers. Values correspond to the range observed across the three interfaces within each assembly. Buried surface area (BSA), number of interface residues, hydrogen bonds, salt bridges and Complex Formation Significance Score (CSS) are shown.

| **Assembly** | **Buried surface area (Å²)** | **Interface residues (per chain)** | **Hydrogen bonds** | **Salt bridges** | **CSS** |
|---|---|---|---|---|---|
| BTV-4 homotrimer | 910-943 | 26-27 | 13-14 | 9 | 0.000 |
| BTV-8 homotrimer | 958-1004 | 22-28 | ~13 | 7 | 0.000 |
| Heterologous trimer<br>(1×BTV-4 + 2×BTV-8) | 873-1074 | 22-32 | 11-15 | 6-7 | 0.000 |
| Heterologous trimer<br>(2×BTV-4 + 1×BTV-8) | 853-971 | 22-28 | 12-15 | 5-9 | 0.000 |

